# Roles of a *Drosophila* ADAM 10 transmembrane metalloprotease, Kuzbanian, in tissue architecture and function of the adult adipose tissue

**DOI:** 10.64898/2025.11.30.691469

**Authors:** Yusaku Hayashi, Taiichi Tsuyama, Tadao Usui, Takumi Ohbayashi, Takefumi Kondo, Kumiko Kokuryoh, Tomonari Awaya, Tatsuya Katsuno, Tadashi Uemura

**Author notes:** **Corresponding authors’ email**. Taiichi Tsuyama: Laboratory of Evolutionary Cell and Developmental Biology, JT Biohistory Research Hall, 1-1 Murasaki-cho, Takatsuki 569-1125, Osaka, Japan Tadashi Uemura: Institute for Life and Medical Sciences, Kyoto University, 53 Shogoin Kawahara-cho, Sakyo-ku, Kyoto 606-8507, Japan.

## Abstract

Various molecular mechanisms in mature adipose tissue regulate whole-body physiology. In contrast, our current understanding of the development of this tissue is limited. The adult fat body (AFB) of *Drosophila melanogaster* is an emerging model for investigating the regulatory mechanisms of adipose tissue morphogenesis in vivo. AFB precursor cells undergo long-range directional and collective migration, proliferation, and homotypic adhesion, leading to the formation of monolayered AFB during metamorphosis. Here, we show that a *Drosophila* ADAM10 transmembrane metalloprotease, Kuzbanian (Kuz), is required for the AFB morphogenesis in the precursor cells at the onset of adhesion. Histological and ultrastructural analyses revealed that the *kuz* knocked-down AFB cells formed multilayered structures or clumps, indicating that Kuz is essential for forming a single-cell-thick tissue sheet. Furthermore, adults with the morphologically altered AFB were supersensitive to starvation stress. Finally, we tested the hypothesis that Notch (N), a well-characterized substrate for Kuz or ADAM10, participates in AFB development, and found that knocking down *N* reduced AFB area coverage and made the adults less adaptive to starvation, similar to *kuz* knockdown. Altogether, our study highlights the critical role of Kuz in both the architecture and function of adult adipose tissue.

## Introduction

Unlike other organs, adipose tissue has a remarkable ability to expand and contract in response to nutritional status in both invertebrates and vertebrates (Gesta et al., 2007; Sakers et al., 2022). Adipocytes store energy metabolites in large lipid droplets upon full feeding, and in the face of starvation, these metabolites are mobilized, contributing to organismal survival. Furthermore, adipose tissue secretes a variety of hormones for the systemic control of metabolism; therefore, it is a critical hub of interorgan communication (Rosen and Spiegelman, 2014; Droujinine and Perrimon, 2016). However, various metabolic disorders unfold once lipids are excessively accumulated, leading to obesity. To prevent such disorders, the pathologies of mature adipose tissue have been extensively studied (Hagberg and Spalding, 2024). In contrast, much less is known about normal developmental processes, including stem cell lineage, tissue morphogenesis, and regulatory genes in vivo, which comprise the basis for investigating the causes of disease states (Gesta et al., 2007; Chi et al. 2018; Sebo and Rodeheffer, 2019; Sakers et al. 2022).

To investigate the genetic programs that ensure adipose tissue development, an emerging in vivo model is the adult adipose tissue of *Drosophila melanogaster*, named adult fat body (AFB). *D. melanogaster* develops two distinct fat bodies during its life cycle: the larval fat body (LFB) and AFB (Figure 1A; Rizki et al. 1980; Li et al., 2019). The LFB develops during embryogenesis and serves as a major energy reservoir in feeding larvae and as a nutriment reserve during metamorphosis and in newly eclosed adult flies (Aguila et al., 2007; Aguila et al., 2013; Boulan et al. 2015; Butterworth et al., 1988; Koyama et al. 2020; Krejčová et al. 2024). By the time of adult emergence, AFB forms anew, completely replaces LFB by 2-3 days after adult eclosion, and plays pivotal roles in systemic controls of metabolism, reproduction, adaptive behaviors, and lifespan until the end of life (Johnson and Butterworth, 1985; Arrese and Soulages, 2010; Storelli et al. 2019). Relatively little was known about how the AFB is formed during metamorphosis (Ferrus and Kankel, 1981; Lawrence and Johnston, 1986; Hoshizaki et al. 1995), until our and others’ in vivo time-lapse imaging of whole-mount pharate adults uncovered many aspects of dynamic behaviors of AFB precursor cells between ∼15 hr after puparium formation (APF) and ∼65 hr APF: long-range directional and collective migration, proliferation, and homotypic adhesion (Figure 1B; Tsuyama et al. 2023 and Lei et al. 2023). Nevertheless, the genes that regulate these dynamic cell behaviors, leading to the formation of monolayered AFB, remain largely undiscovered, as does the functional relevance of this tissue architecture.

**FIGURE 1.**
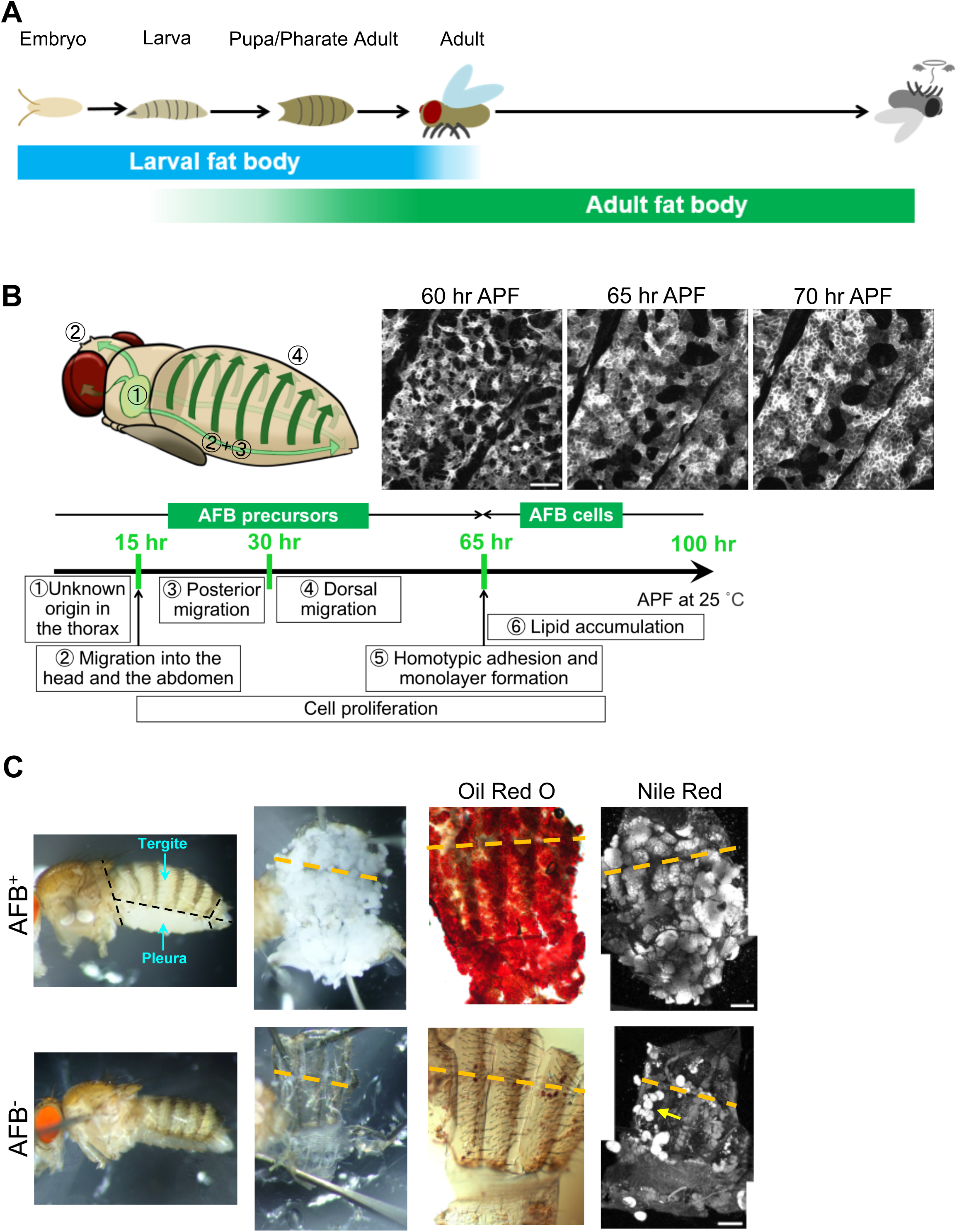
Development of adult fat body (AFB) in *Drosophila melanogaster* and its defective formation observed in a wildtype strain. (A) Two distinct fat bodies in the life cycle of *Drosophila melanogaster*: larval fat body (LFB) and adult fat body (AFB). The LFB persists until 2-3 days after adult eclosion and is completely replaced by the AFB, which is formed de novo in pupae and pharate adults. (B) Schematic illustration of the migration pathways of AFB precursors (upper left) and timeline of AFB development during metamorphosis (bottom; adapted from Tsuyama et al. 2023). We refer to cells prior to the AFB formation (approximately 65 h APF) as AFB precursor cells (the precursor cells for short) and cells after the formation as AFB cells. For simplicity, the pathways are drawn in a pharate adult and numbered as in the timeline, whose scales represent hours after puparium formation (APF) at 25°C. Adults emerge approximately 96 hr APF. AFB precursors and AFB cells in this timeline can be visualized using *c833-Gal4+ppl-4kb-Gal80* (Tsuyama et al. 2023) or *OK6-Gal4* (Lei et al. 2023). These driver strains were also used to knock down genes in this study (see details in “Fly strains and husbandry” in Experimental Procedures). (Upper right) Time-lapse images of the precursors labeled with *OK6-Gal4>CD4::tdGFP* in a whole-mount pharate adult. The precursors underwent homotypic adhesion and monolayer formation (circled number 5 in the timeline). The anterior is to the top left and the dorsal is to the top right. (C) A wild-type strain, *Canton-S*, showed disorganized or lost AFB tissues with approximately 30% penetrance (Tsuyama et al., 2023; see also Figure S1A). Representative images of 7-8-day female adults with fully developed AFB (AFB^+^, top row) or underdeveloped AFB (AFB^-^, bottom row). Abdomens were dissected along the black dashed lines shown in the far-left image, pinned down (second from the left), and stained with Oil Red O (second from the right) or Nile Red (far right). The orange dashed lines indicate the dorsal midline. The arrow marks an example of a cluster of remaining LFB cells. n (Oil Red O)=3 (AFB^+^) and 5 (AFB^-^), and n (Nile Red)=10 for each group. Scale bars: 50 µm in B and 200 µm in C.

We previously reported that wildtype strains exhibit aberrant AFB development with variable penetrance (Tsuyama et al., 2023), which might be due to genetic variations. Here, we took advantage of this phenotypic variability among the wild-type strains to identify and characterize novel candidates for genes controlling AFB development. We performed a targeted RNAi screen of the candidates and found that knocking down *kuzbanian* (*kuz*), which encodes a homolog of the ADAM10 transmembrane metalloprotease, in AFB precursor cells significantly reduced the area coverage of AFB. To scrutinize the *kuz* knocked-down tissue architecture, we first identified AFB soon after formation in control pharate adults under light and electron microscopes and then characterized how the architecture of the young AFB was affected by the *kuz* knockdown. Furthermore, we addressed whether a transmembrane protein, Notch (N), which is one of the best characterized substrates for Kuz or ADAM10, participates in the AFB development. Finally, we addressed whether the detected morphologically abnormal AFB was fully functional in adults. Together, our results define a critical role for the ADAM10 metalloprotease Kuz in sculpting the architecture of adult adipose tissue and suggest that this developmental process is essential for its metabolic function.

## Results

### A *Drosophila* ADAM10 transmembrane metalloprotease, Kuzbanian (Kuz), is a candidate regulator of AFB development

AFB predominantly resides in the abdomen, which appears white and massive in dissected adults, and the stored neutral lipids in the droplets are stained with Oil Red O or Nile Red (see an example in “AFB^+^” in Figure 1C). Adults of *Canton-S*, a widely used wildtype strain, unexpectedly showed disorganized or lost AFB tissues with approximately 30% penetrance (see a severe example in “AFB^-^” in Figure 1C). Moreover, representative 39 strains in *Drosophila* Genetic Reference Panel (DGRP; Huang et al., 2014; MacKay et al., 2012) also showed such a phenotype with variable degrees of penetrance below 12% (Tsuyama et al., 2023). This variation allowed us to perform a genome-wide association (GWA) analysis to explore the genetic variants associated with defective AFB development (Figure S1A, B). We identified a total of 148 single nucleotide polymorphisms (SNPs) in or near 88 genes, of which we selected 15 candidate genes based on GO terms related to “stem cell regulation,” “cell migration,” and/or “cell adhesion” (Table S1).

We subsequently examined whether knocking down each candidate gene affected AFB development. For this purpose, we expressed hairpin RNAs in AFB precursor cells and observed their effects on AFB in whole-mount pharate adults shortly before eclosion (Figure S1C; Table S1; see the exact genotypes and the numbers of flies observed of this and subsequent experiments in Table S2 and individual figure legends, respectively). The list of the candidate genes included the serpent (*srp*) gene, which encodes a GATA-type transcription factor (Hayes et al., 2001). *srp* served as a positive control for this screen because its requirement for the AFB development was previously reported (Lei et al., 2023; Tsuyama et al., 2023) and its knockdown in the precursors caused almost complete loss of the *CD4::tdGFP*-positive AFB cells (“*srp* KD” in Figure S1C). The candidate gene list includes *heartless* (*htl*), which encodes the FGF receptor required for the directional migration and spreading of AFB precursors (Lei et al., 2023), and its knockdown also severely affected AFB formation (“*htl* KD” in Figure S1C).

Among the other candidate genes, knocking down *kuzbanian* (*kuz*) appeared to reduce AFB area size, whose defect was milder than that of the *srp* or *htl* knockdown (“*kuz* KD” in Figure S1C), while knocking down any of the remaining 12 genes did not give rise to an obvious phenotype. We further examined the effect of the *kuz* RNAi construct on AFB in dissected abdomens of pharate adults by staining with Nile Red (Figure 2A). A wide belt of AFB was formed in each abdominal segment in the control fly, whereas the belt became narrower and the Nile Red signal-free areas expanded in the *kuz* KD fly, in particular, near the dorsal midline (towards the top of the panels in Figure 2A), suggesting that *kuz* is required for AFB development.

**FIGURE 2.**
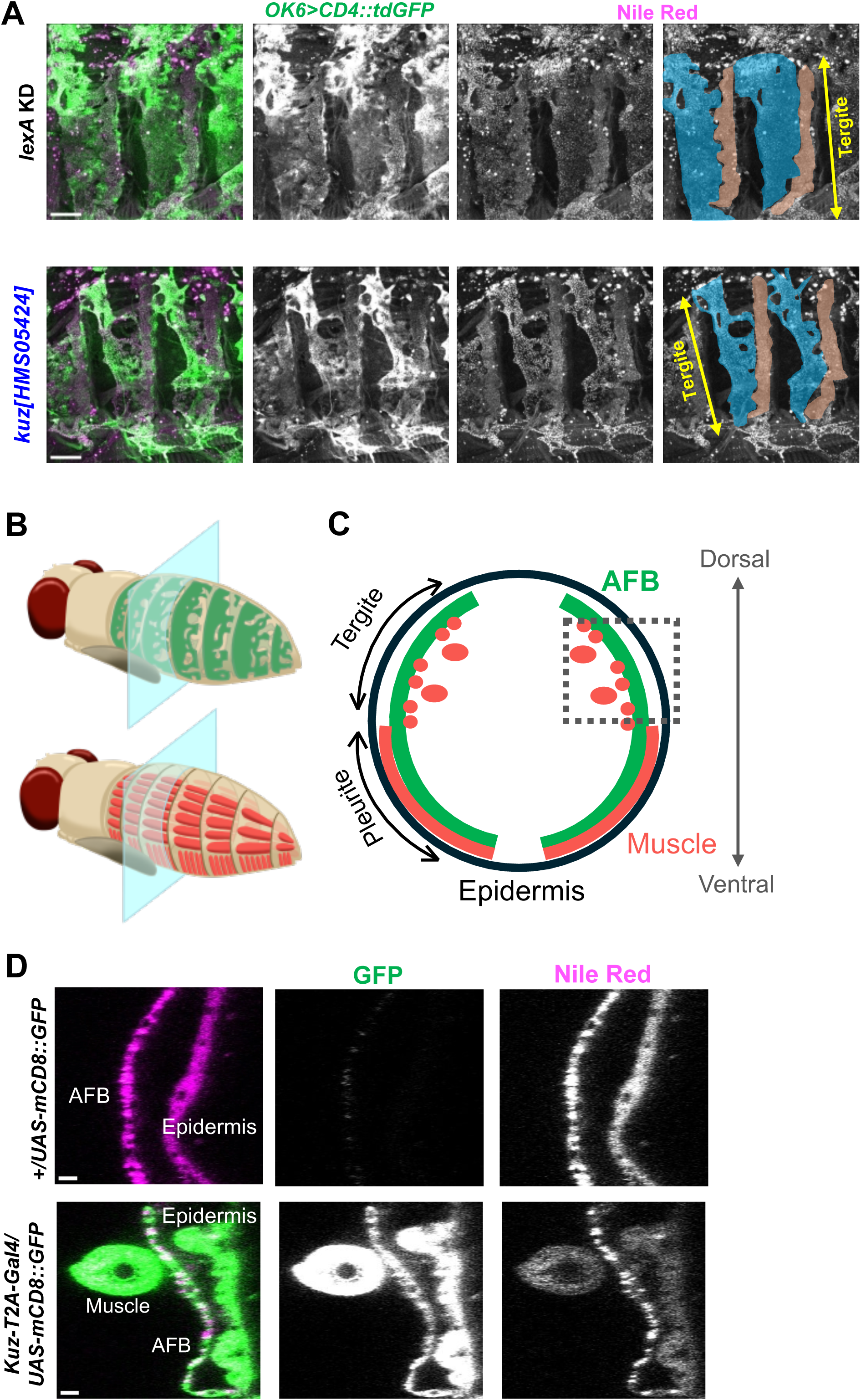
Reduction in the AFB area size in pharate adults when *kuzbanian* (*kuz*) was knocked down and *kuz* expression patterns in the wildtype pharate adult. (A) Representative images of dissected abdomens of pharate adults (73 hr APF at 25°C). (Top row) Control pharate adult. The control short hairpin RNA for the knockdown (KD) was *lexA*. (Bottom row) *kuz* knockdown pharate adult using *OK6-Gal4* and *kuz[HMS05424]*. *CD4::tdGFP* signals were detected using an anti-GFP antibody, and lipid droplets were stained with Nile Red. In the far-right panel, AFB and oenocytes are colored blue and brown, respectively. In all images, the anterior is to the left and the dorsal is upwards. n=3 for each group. Scale bars: 100 µm. (B, C) Schematic illustrations of AFB (green) and muscles (orange) in the abdomens of pharate adults (B). Transverse sections of the abdomen (semitransparent cyan boards in B) are enlarged in “C,” which indicates the spatial relationships of the AFB, muscles, and epidermis. In the tergite (dorsal region), the AFB is located between the epidermis and muscles, whereas the AFB in the pleurite (ventral to the tergite) is the innermost of the three tissues (Tsuyama et al. 2023). (D) Reconstructed images of the three tissues in the transverse section (within a part of the gray dashed box in “C”) with the body cavity to the left. These images were made using those taken from dissected abdomens at 73 hr APF, which were stained with Nile Red. (Top row) Negative control pharate adult that did not express GFP in any of the tissues. (Bottom row) Pharate adult of *kuz-T2A-Gal4>CD4::tdGFP* in which *kuz*-expressing tissues emit green fluorescence. Note that the GFP signals of the muscles and epidermis were over-intensified to detect the AFB signal. n=2 (top) and 3 (bottom). Scale bars: 10 µm.

In the *kuz* gene, we identified a total of 6 genetic variants that are significantly associated with the defective AFB development, all of which are located in the 2.06 kb genomic region that includes the third exon (Figure S1D and its legend; “2L_13558032_DEL”∼“2L_13560082_SNP” in Table S3). In each of the 39 DGRP strains in which we analyzed AFB, those 6 variants were either all minor alleles or all major ones, and three strains out of the top four strains with high penetrance of the defect possess the minor set (Table S3).

*kuz* encodes a *Drosophila* homolog of the ADAM10 transmembrane metalloprotease that contains an extracellular protease domain and regulates the molecular functions of diverse substrate proteins by shedding their ectodomains (Weber and Saftig, 2012; Lambrecht et al. 2018; Rosenbaum and Saftig 2024). *kuz* is an essential gene for viability, expressed at high levels in the nervous system and imaginal discs in larvae, and plays pivotal roles in neurogenesis and many other developmental contexts, including collective cell migration (Sotillos et al. 1997; Fambrough et al. 1996; Rooke et al. 1996; Pan and Rubin 1997; Wang et al. 2006; Brown et al., 2014).

### *kuz* is expressed in AFB

To find evidence of *kuz* expression in AFB precursor cells and AFB cells (see the definitions in the Figure 1B legend) in pharate adults, we addressed whether *kuz* is transcribed in these cells using a *kuz-T2A-Gal4* stock (Diao and White, 2012). In the genome of this stock, a *T2A-Gal4* cassette is inserted in the intron of *kuz*, which is located between the exons coding the prodomain. This *kuz-T2A-Gal4* stock was crossed with a *UAS-mCD8::GFP* stock (Lee and Luo, 1999). 73 hr APF pharate adults of the next generation were collected, their abdomens were dissected and stained with Nile Red; and acquired images were reconstructed on the transverse section (Figure 2B, C, D). AFB was positive for GFP, although the signal was weaker than that of the epidermis and muscle (“*Kuz-T2A-Gal4/UAS-mCD8::GFP*” in Figure 2D), and none of these GFP signals of the three tissues was autofluorescence because it was not seen in the absence of *Kuz-T2A-Gal4* (“*UAS-mCD8::GFP*” in Figure 2D). Although *kuz* expression in pharate adults has not been studied by immunostaining or RNA in situ hybridization, *kuz* is transcribed in a wide range of larval and adult tissues (Sotillos et al. 1997; Brown et al. 2014). Together, our results strongly suggest that *kuz* is transcribed in the AFB in pharate adults. Although we found GFP-positive migrating cells in the whole-mount pharate adult at 50 hr APF, it was difficult to conclude whether they were AFB precursor cells because of the strong epidermal signals that obscured the precursors or other motile cell types, such as hemocytes (data not shown).

In addition to our results indicating *kuz* transcription in the AFB, we attempted to visualize GFP-tagged Kuz protein during the AFB development by using a previously reported “GFP-tagged Kuz in BAC” stock (Dornier et al., 2012) or by generating a new one with the help of an available MiMIC stock (the “MiMIC GFP-tagged Kuz” stock; Figure S2; Nagarkar-Jaiswal et al., 2015). Although we detected GFP signals in larval central nervous system and imaginal discs in our MiMIC GFP-tagged Kuz stock, either attempt to detect GFP signals in AFB was unsuccessful likely due to low expression levels (see details in the Figure S2 legend and Figure 3E below).

**FIGURE 3.**
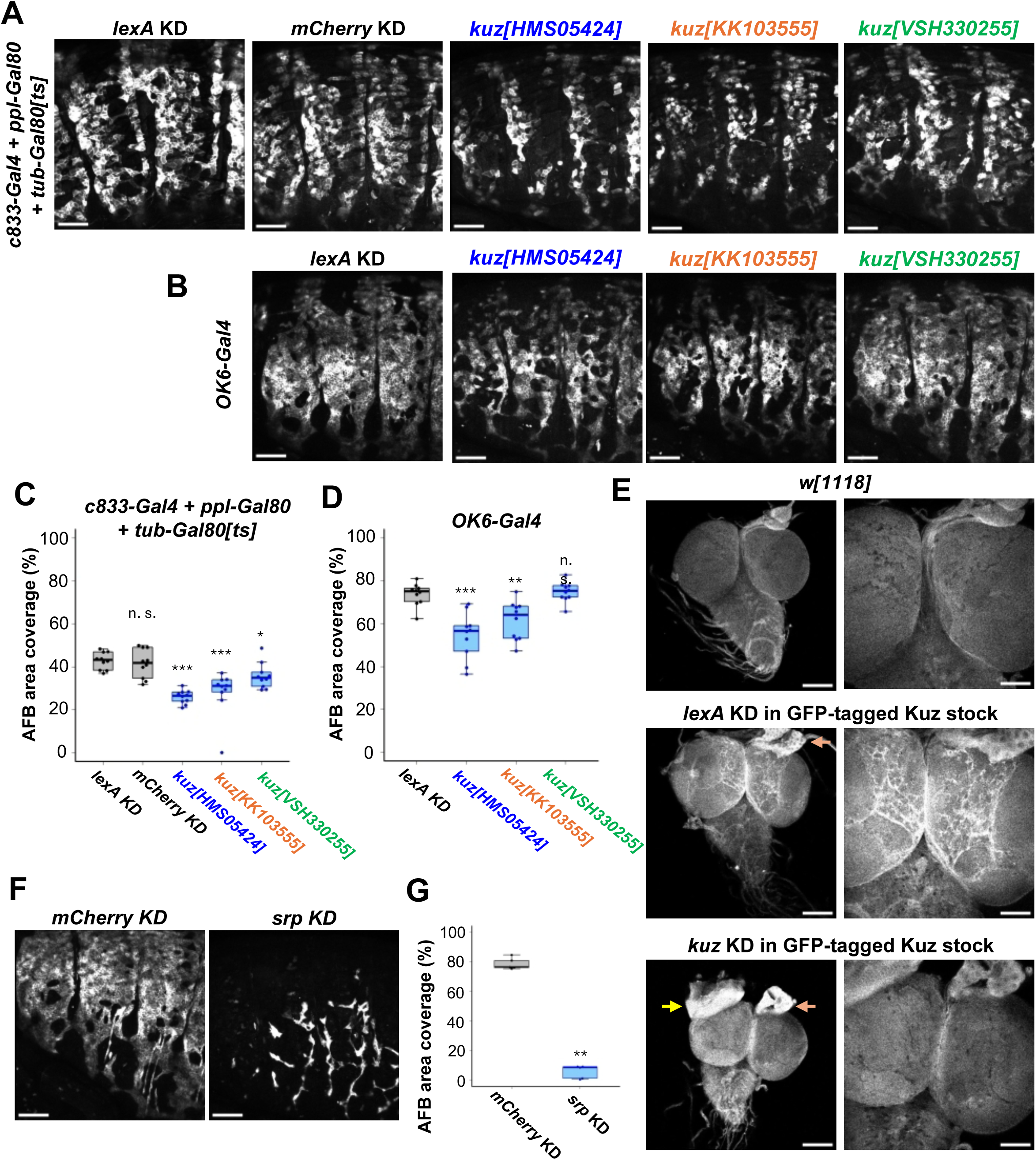
Quantifications of the AFB area coverage in pharate adults when *kuzbanian* (*kuz*) or *serpent* (*srp*) was knocked down. (A, B) Lateral views of the abdomens of whole-mount pharate adults where *kuz* was knocked down and AFB cells were visualized with *CD4::tdGFP* that was expressed by either *c833-Gal4+ppl-4kb-Gal80+tub-Gal80[ts]* (A) or *OK6-Gal4* (B). The genomic locations of the three target sequences are shown in Figure S1D. Control short hairpin RNA for knockdown (KD) were *lexA* or *mCherry*. (A) Flies were aged at 19°C to inhibit *Gal4* expression and shifted to 29°C (see details in “Fly strains and husbandry” in Experimental Procedures). Images were captured at 73 hr APF at 29°C, which was equivalent to a stage shortly before eclosion. (B) The flies were kept at 25°C throughout the experiment, and images were captured at 73 hr APF. Compared to the image of the control fly in “B,” *CD4::tdGFP* signals in the controls in “A” appeared patchy. This is most likely due to the onset of *ppl-4kb-Gal80* expression in maturing AFB cells. The purpose of establishing the *ppl-4kb-Gal80* stock was to conceal GFP expression in fully matured larval fat body cells that remain during metamorphosis (Tsuyama et al. 2023). (C, D) Quantification of AFB area coverage in pharate adults of the genotypes in “A” and “B” (C and D, respectively). n=10 (A and C) and 10 (B and D). (E) Dorsal views of the entire central nervous system of wandering larvae (left column) and enlarged images of the brains (right column). (Top) Control strain, *w[1118]*, which does not express GFP. n=2. (Middle and bottom) GFP-tagged Kuz stock (Figure S3) that expresses GFP in the cortex glia. In contrast to the negative control for the knockdown (middle), the GFP signal of the cortex glia was lost in the *kuz* knockdown using *Repo-Gal4* (bottom). Yellow and orange arrows indicate the eye-antenna disc and ring glands, respectively. n=4 (middle) and 4 (bottom). (F, G) Lateral views of the abdomens of whole-mount pharate adults at 73 hr APF, where *serpent* (*srp*) was knocked down using *OK6-Gal4* (F) and quantification of the AFB area coverage (G). n=5. In all images of the lateral view, the anterior is to the left and the dorsal is upwards. Steel’s test. *p<0.05, **p<0.01, and ***p<0.001. n.s.: not significant. Scale bars: 100 µm in (A), (B), (E) (left column), and (F); 50 µm in (E) (right column). To knock down *kuz* in AFB development in subsequent experiments, *OK6-Gal4, kuz[HMS05424]*, and *lexA* (a negative control) were employed unless described otherwise.

### *kuz* is a novel regulatory gene for AFB development

In the aforementioned RNAi screen of the candidate genes, we employed a single RNAi line (*kuz[HMS05424]*) to knock down *kuz*. To minimize the concern of an off-target effect, we employed two additional RNAi lines whose target sequences partially overlap (kuz[KK103555] and kuz[VSH330255]; Figure S1D and its legend; Öztürk-Çolak et al., 2024). Individual hairpin RNAs were expressed in AFB precursor cells and AFB cells during metamorphosis by two distinct sets of drivers: *Gal4+ppl-4kb-Gal80+tub-Gal80[ts]* and *OK6-Gal4* (Figure 3A, B; see details in “Fly strains and husbandry” in Experimental Procedures). The effects of RNA expression on AFB area coverage were quantitatively assessed (Figure 3C, D; see details in Experimental Procedures). Compared to the controls, all three RNAs, or two of the three, *kuz[HMS05424]* and *kuz[KK103555]* that do not share the target sequence, significantly reduced the area coverage (Figure 3C, D). Among the three RNAs, *kuz[HMS05424]* was the most effective (Figure 3C, D). Taken together, the similar defects in AFB coverage were caused by expressing at least two distinct RNAs using two different driver sets, suggesting that the aberrant AFB developmental phenotype was due to Kuz depletion and not an off-target effect. As previously reported, knocking down *srp* caused a much more severe phenotype (Figure 3F, G).

We examined the knockdown efficacy of *kuz[HMS05424]* using the MiMIC GFP-tagged Kuz stock we generated (Figure S2). Expression of *kuz[HMS05424]* selectively in glial cells caused the GFP signal in the cortex glia to almost disappear, while the signals in the eye imaginal disc and the ring gland remained (Figure 3E), indicating that the expression of this RNA reduced the amount of Kuz efficiently. Taken together, our results strongly suggest that *kuz* is required for AFB development.

### *kuz* contributes to AFB development in the stages of the precursor cell adhesion and monolayer formation

To further investigate how *kuz* regulates AFB development temporally and structurally, we employed the following stocks in our subsequent experiments: *kuz[HMS05424]* as the RNAi stock, *OK6-Gal4* as the driver to make the fly genotypes simpler, and *lexA[HMS05772]* as the negative control that shares the same landing site as *kuz[HMS05424]*.

We first addressed the step of AFB development for which *kuz* is required. For this purpose, we traced back to the earlier stages of AFB development (60-70 hr APF). During these stages after the migration period, the precursor cells still extended prominent filopodia-like protrusions, increased contact areas with neighboring cells as fine protrusions were retracted, and eventually formed the characteristic monolayer, as highlighted in Figure 1B (Tsuyama et al., 2023; Figure 4). As shown in our quantification of the AFB area coverage, there was no significant difference between the control and *kuz* knockdown pharate adults before the adhesion of the precursor cells (60 hr APF). In contrast, at the onset of adhesion (65 hr APF) and soon after adhesion (70 hr APF), the area coverage was significantly decreased in the *kuz* knockdown compared to the control (Figure 4A, B). We further tracked the coverage dynamics of individual pharate adults along the timeline (Figure 4C). The coverage increased between 60 and 65 hr APF in the control, whereas it increased only slightly in the *kuz* knockdown between these time points. In conclusion, *kuz* is required in the stages of precursor cell adhesion and monolayer formation to expand the AFB to cover the abdomen almost entirely.

**FIGURE 4.**
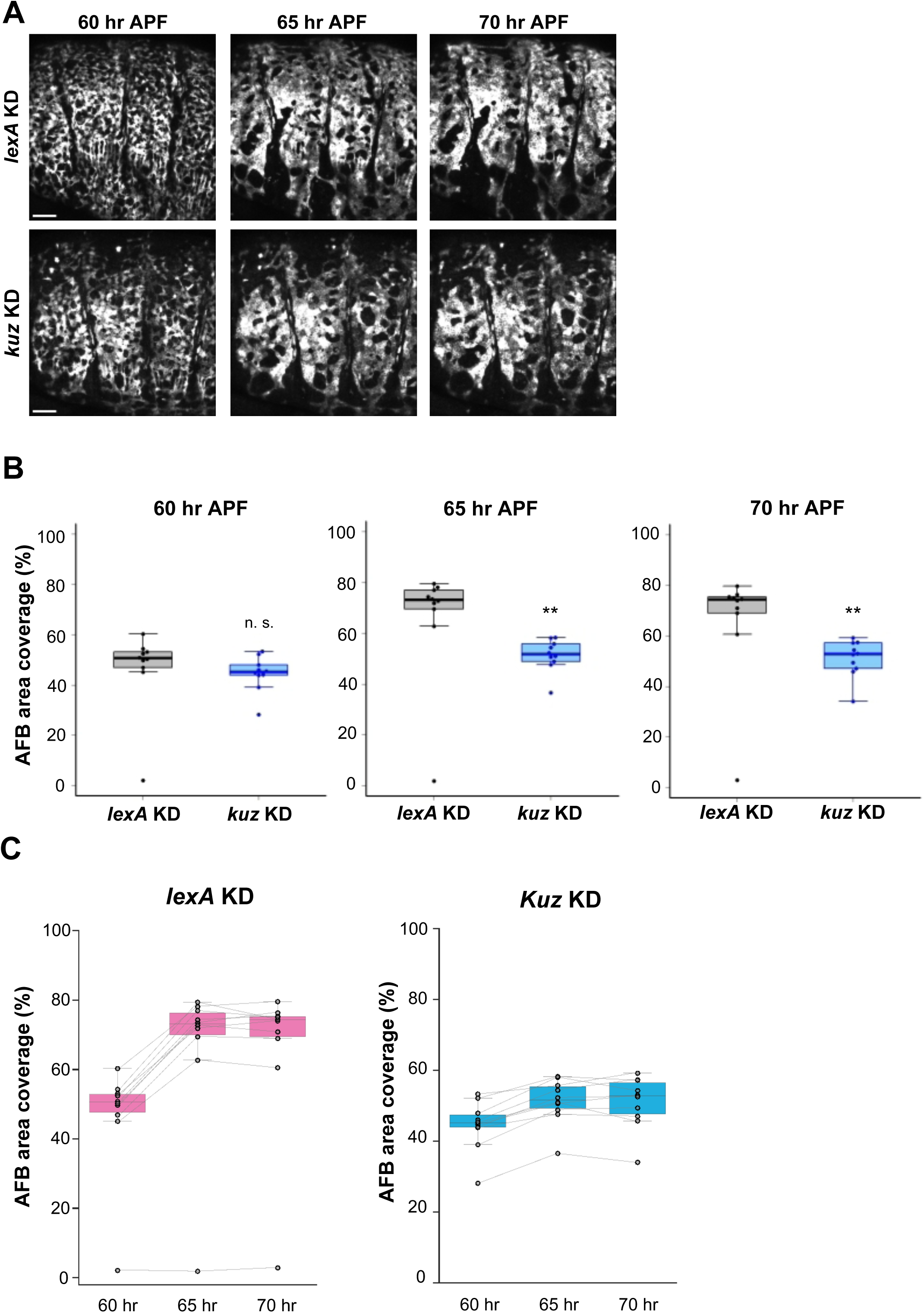
Effects of the *kuz* knockdown at the stages of precursor cell adhesion and monolayer formation. (A) Time-lapse images of pharate adults of the control (top row, *lexA* KD) and the *kuz* knockdown (bottom row, *kuz* KD) during precursor cell adhesion and monolayer formation. In all images, the anterior is to the left and the dorsal is upwards. (B) Quantification of AFB area coverage at individual time points. *p<0.05 and **p<0.01. Wilcoxon-Mann-Whitney test. Scale bars: 100 µm. (C) The datasets in “B” were used to show time series variations in AFB area coverage of individual pharate adults of the control (*lexA* KD) or the *kuz* knockdown genotype (*kuz* KD). n=10 for each time point of each genotype.

### Histological analysis showed that AFB became fragmentary in the *kuz* knockdown pharate adult, in contrast to the continuous layer in the control

How is the AFB area coverage reduced in *kuz* knockdown pharate adults? To dissect the cellular basis of this phenotype, it was crucial to observe developing AFBs from surface to depth under light and electron microscopes. To achieve this goal, we first identified the tissue in the abdominal sections of the control pharate adults and then investigated how this tissue morphology was affected by *kuz* knockdown. Below, we conducted anatomical analyses of flies at 73 hr APF.

We first aimed to identify the AFB in the frontal sections at the dorsal level of the control pharate adult (Figure 5A, B). In each abdominal segment, the AFB is located anterior to the pheromone-producing organ, oenocytes (see the far-right column of Figure 2A; Makki et al., 2014), and our previous whole-mount imaging showed that the AFB is located between the epidermis and muscles at the dorsal level (see those tissues’ relative position in “Tergite” in Figure 2C; Tsuyama et al., 2023). This spatial information of the surrounding tissues allows us to assume that a thin layer extending along the anterior-posterior axis (the A-P axis) underneath the epidermis is the young AFB (Figure 5C, green in Figure 5D). Consistent with our assumption, the equivalent tissue in the genotype expressing *CD4::tdGFP* in AFB was positive for GFP (Figure S3). Moreover, such a thin tissue was missing in the *srp* knockdown (Figure 5E, F), as expected from the almost complete loss of AFB caused by knocking down *srp* (Figure S1C). These results support our assumption that the thin layer is a young AFB. In the *kuz* knockdown pharate adults, the epidermis, muscles, and oenocytes were formed as in the control, and thin tissues were aligned along the A-P axis between the epidermis and muscles; however, these thin tissues were shorter and discontinuous compared to the AFB in the control (Figure 5G, H; Figure S4A-C).

**FIGURE 5.**
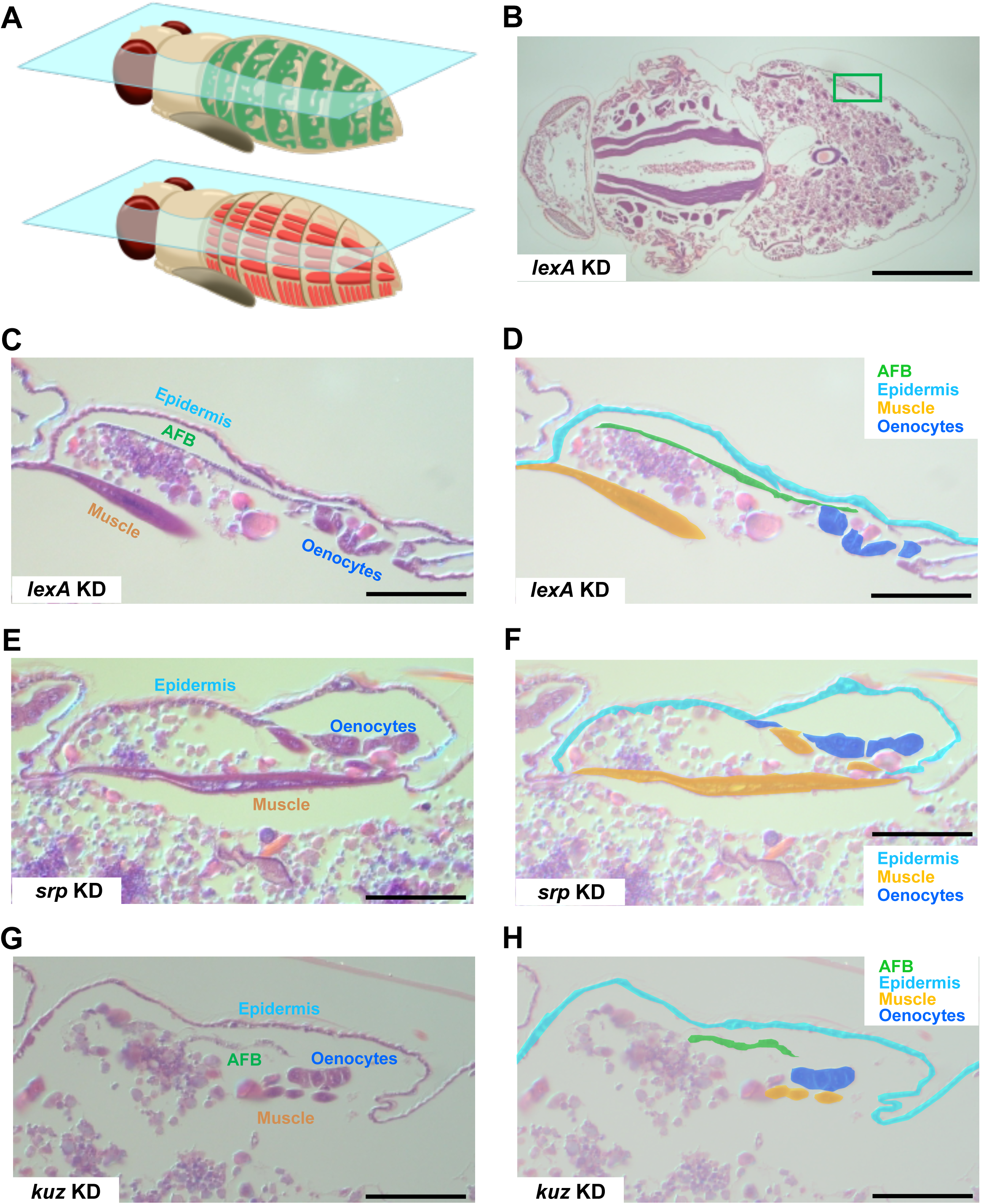
Histological analysis of AFB and the surrounding tissues at the dorsal level. (A) Schematic illustrations of AFB (green) and muscles (orange) in the abdomens of pharate adults and frontal sections at the dorsal level (semitransparent cyan boards). (B-H) Representative images of the frontal sections at 73 hr APF of the control (B, C, D), *srp* knockdown (E, F), and *kuz* knockdown (G, H). The body surface area of the abdomen of the control (green box in B) is enlarged, and the identified tissues are labeled (C) or pseudocolored to indicate individual tissues (D). Similarly, images of the corresponding regions of the *srp* or *kuz* knockdown are shown in (E, F, G, H). In “D,” “F,” and “H,” the brightness of the original images (C, E, and G, respectively) was increased to highlight the coat. See the spatial relationships between AFB, muscles, and epidermis in the dorsal region (tergite) in Figure 2C. In all images, anterior is to the left. Scale bars: 500 µm in B; 50 µm in C-H. n (microscope fields)=20 (*lexA* KD), 18 (*srp* KD), and 15 (*kuz* KD) including the one in Figure S4C. n (flies)=5 (*lexA* KD), 3 (*srp* KD), and 3 (*kuz* KD).

We similarly examined frontal sections at the ventral level of the control pharate adult (Figure 6A, B), where the AFB is located more internally than the muscles, in contrast to its position relative to the muscles at the dorsal level (see “Pleurite” in Figure 2C; Tsuyama et al., 2023). We identified AFB located deeper than the muscle bundles that run along the dorsal-ventral axis (Figure 6C, green in Figure 6D). As seen in the sections at the dorsal level, AFB was missing in the *srp* KD (Figure 6E, F), and in the *kuz* knockdown, we found short and discontinuous thin tissues (Figure 6G, H; Figure S4D-F). At both dorsal and ventral levels, this morphological feature is consistent with our observations of the narrower AFB along the A-P axis and the smaller coverage of the AFB in the *kuz* knockdown pharate adults (Figure 2A and Figure 3A-D, respectively), indicating that these tissues are fragmentary AFB.

**FIGURE 6.**
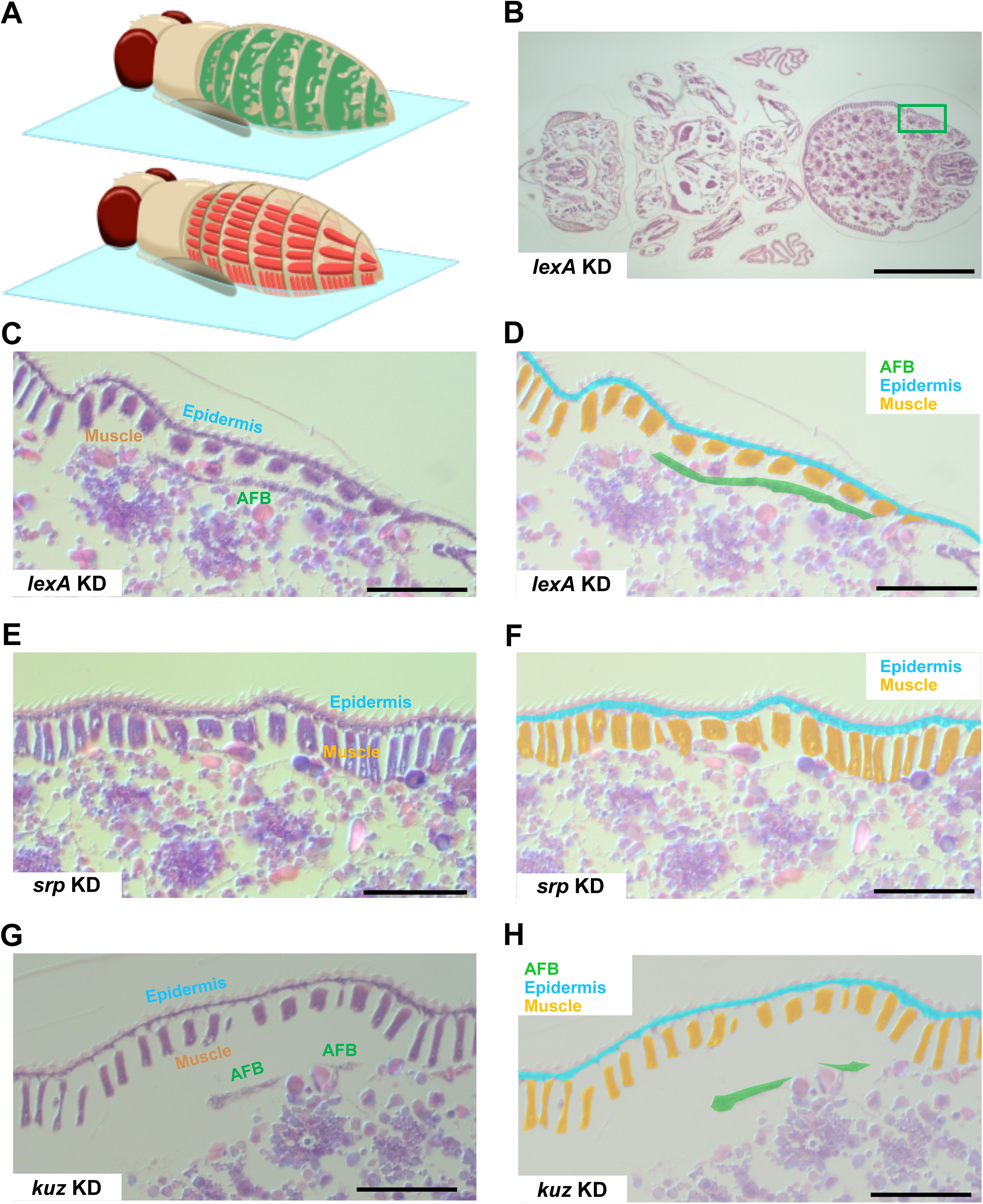
Histological analysis of AFB and the surrounding tissues at the ventral level. (A) Schematic illustrations of AFB (green) and muscles (orange) in the abdomens of pharate adults and frontal sections at the ventral level (semitransparent cyan boards). (B-H) Representative images of the frontal sections at 73 hr APF of the control (B, C, D), *srp* knockdown (E, F), and *kuz* knockdown (G, H). The body surface area of the abdomen of the control (green box in B) is enlarged, and the identified tissues are labeled (C) or pseudocolored to indicate individual tissues (D). Similarly, images of the corresponding regions of the *srp* or *kuz* knockdown are shown in (E, F, G, H). In “D,” “F,” and “H,” the brightness of the original images (C, E, and G, respectively) was increased to highlight the coat. See the spatial relationships of AFB, muscles, and epidermis in the ventral region (pleurite) in Figure 2C. In all images, anterior is to the left. Scale bars: 500 µm in B; 50 µm in C-H. n (microscope fields)=27 (*lexA* KD), 12 (*srp* KD), and 19 (*kuz* KD) including the one in Figure S4G. n (flies)=5 (*lexA* KD), 3 (*srp* KD), and 3 (*kuz* KD).

### Ultrastructural analysis revealed that the *kuz* knockdown AFB cells formed either multi-cell-layered structures or clumps

To compare the AFB of the two genotypes at the cellular level, we observed ultrathin sections of pharate adults under an electron microscope. In our observations of the frontal sections at the dorsal level, we first identified a thin layer of AFB in the control flies based on the relative positions of the individual tissues, as we did in the light microscopic analysis (Figures 7A, B). Higher power images clearly visualized the monolayered structure of AFB and lipid droplets that AFB cells had already accumulated after monolayer formation (Figures 7C-F; Tsuyama et al. 2023; Lei et al. 2023).

**FIGURE 7.**
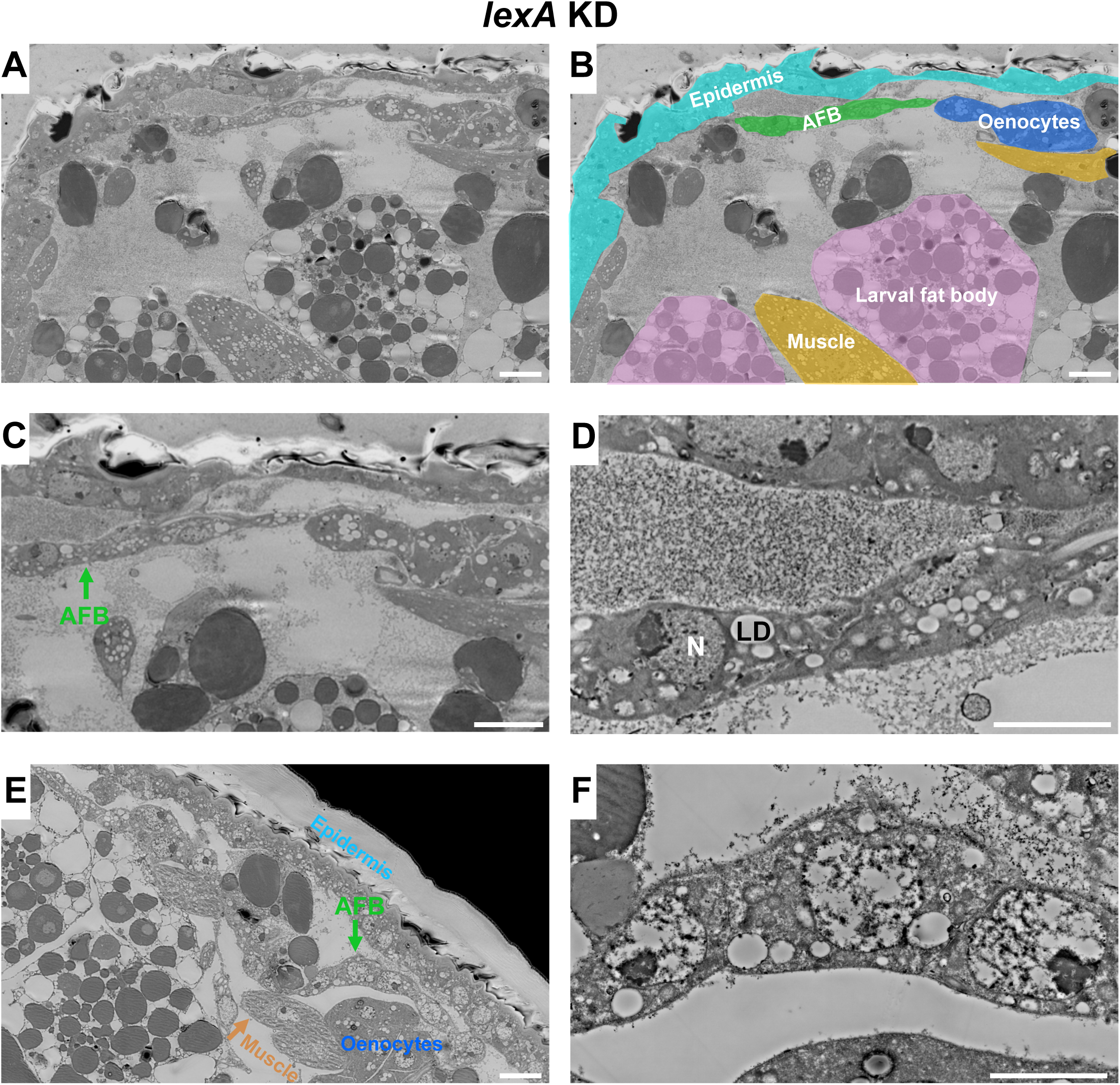
Ultrastructural analysis of AFB in the control pharate adults. Representative backscattered electron (BSE) images of the frontal sections at the dorsal level of the abdomen of the control at 73 hr APF. In the abdomen, these sections were located similarly to that shown in Figure 5C. (A, B) The identified tissues in “A” are labeled and pseudocolored in “B”. Larval fat body cells include electron-dense bodies, such as protein granules (Butterworth et al. 1988; Krejčová et al. 2024). (C, D) AFB in “A” is zoomed in at two distinct magnifications. (E, F) Images taken from a different pharate adult. The magnification of “E” is equal to that of “A, B” and the ratio of “F” is equal to that of “D.” N, nucleus; LD, lipid droplet. In all images, anterior is to the left. Scale bars: 10 µm in A, C, and E; 5 µm in D and F. n (microscope fields)=5 and n (flies)=2. “A”-“D” and “E”-“F” were taken from the different pharate adults.

In the *kuz* knockdown pharate adults, we found tissues with unusual morphologies underneath the epidermis (Figure 8): a cluster of cells in a kinked shape (Figure 8A-D), a multi-cell-layered tissue (Figure 8E, F), and a clump of cells (Figure 8G, H). Although tissues with such overall forms were not observed in the control, all these cells contained lipid droplets, indicating that the individual cells were differentiated adipocytes and that these tissues were indeed AFBs (Figures 8D, F, and H). These abnormal morphologies of AFB were found at the dorsal edge of the tergite (Figure 2C); however, the AFB layer spread in the lateral-to-ventral region (“Plurite” in Figure 2C) of the *kuz* knockdown pharate adults as in that of the control (Figure S5).

**FIGURE 8.**
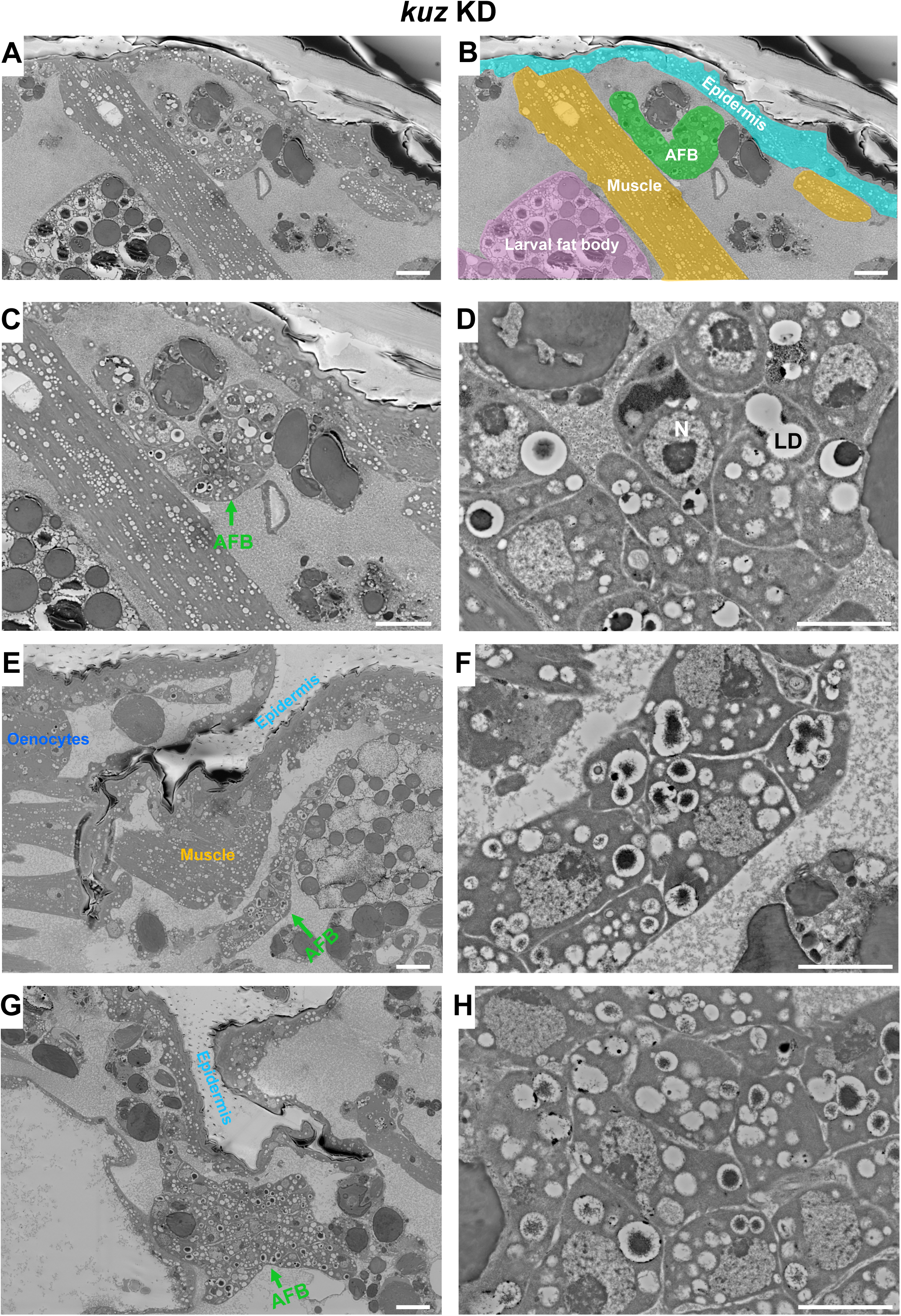
Ultrastructural analysis of AFB in the *kuz* knockdown pharate adults. Representative BSE images of the frontal sections at the dorsal level of the abdomen of *kuz* knockdown larvae at 73 hr APF. In the abdomen, these sections were located similarly to that shown in Figure 5C. (A, B) The identified tissues in “A” are labeled and pseudocolored in “B”. (C, D) AFB in “A” is zoomed in at two distinct magnifications. (E, F) and (G, H) The magnification of “E, G” is equal to that of “A, B” and the ratio of “F, H” is equal to that of “D.” N: nucleus; LD: lipid droplet. Lipid droplets usually become empty during sample preparation for electron microscopic observations, but sometimes residual lipids are seen in the droplets in “D,” “F,” and “H.” In all images, the anterior is to the left. Scale bars: 10 µm in (A), (C), (E), and (G); 5 µm in (D), (F), and (H). n (microscope fields)=8 and n (flies)=2. “A”-“D” and “E”-“F” were taken from the different pharate adults.

We suspected that the abnormal morphogenesis of AFB in the *kuz* knockdown pharate adults might be due to malformation of extracellular matrix (ECM) that is assumed to ensheathe the zone-cell-thick AFB. To visualize ECM in pharate adults (73 hr APF) and newly eclosed adults (day 0-1), we used a Viking (Vkg)-GFP stock, a GFP-trap line of the α2 subunit gene of collagen IV (Morin et al., 2001). At the limit of our observation, we detected the GFP signal that enclosed the AFB layer in both the control and *kuz* knockdown pharate adults (arrows in Figure S6), suggesting that ECM formation was not severely impaired by knocking down *kuz*.

### Lipid storage and tissue morphology of the *kuz* knockdown AFB in adults

In all above experiments using *OK6-Gal4*, the hairpin RNA was expressed in the AFB precursor cells and in the AFB cells after the monolayer formation, while the RNA expression was decreased shortly before eclosion (Figure S7). Because AFB plays pivotal roles in systemic controls of metabolism throughout the adult stage, we examined the effects of such a transient knockdown on the tissue morphology and function of AFB in adults (Figure 9). For simplicity, we designated AFB in adults that experienced *kuz* knockdown during metamorphosis and such adults as the *kuz* knockdown AFB in adults and the *kuz* knockdown adults, respectively.

**FIGURE 9.**
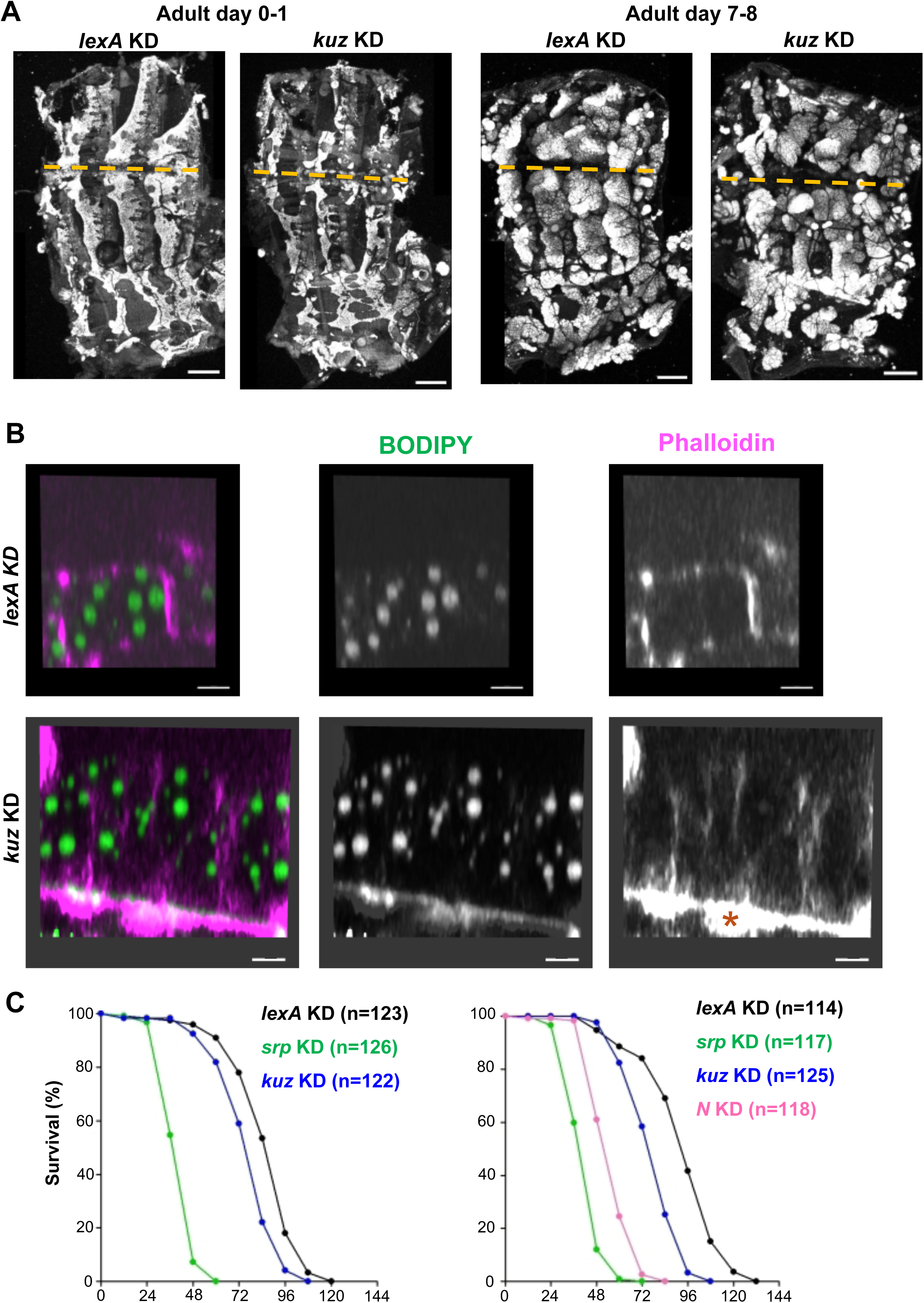
Effects of the *kuz* knockdown on lipid storage, tissue morphology of AFB, and sensitivity to starvation in the adult stages. (A) Abdomens of control or *kuz* knockdown adults were dissected at 0-1 or 7-8 days after eclosion, and neutral lipids were stained with Nile Red. The orange dashed lines indicate the dorsal midline of individual adults. In all images, anterior is to the left. n=4 (*lexA* KD of 0-1 days), 4 (*kuz* KD of 0-1 days), 3 (*lexA* KD of 7-8 days), and 4 (*kuz* KD of 7-8 days). Scale bars: 200 µm. (B) Images corresponding to the portions of the transverse section located near the dorsal midline. The midline is to the left, and the epidermis is downwards. See also the illustration of the section in Figure 2C. Abdomens of control or *kuz* knockdown adults (0-4 hr after eclosion) were dissected, and neutral lipids and cell cortex were stained with BODIPY and Phalloidin 647, respectively. Images were captured using a two-photon microscope. Asterisks indicate autofluorescence. n=3 (*lexA* KD) and 4 (*kuz* KD). Scale bars, 10 µm. (C) Survival curves of *lexA* (control), *srp*, *kuz*, or *N* knockdown adults at 2.5-3 days after eclosion under starvation (n = 114-126). The results of two independent experiments are shown. In all combinations, the differences were significant (p<0.000001) in the log-rank tests.

**FIGURE 10.**
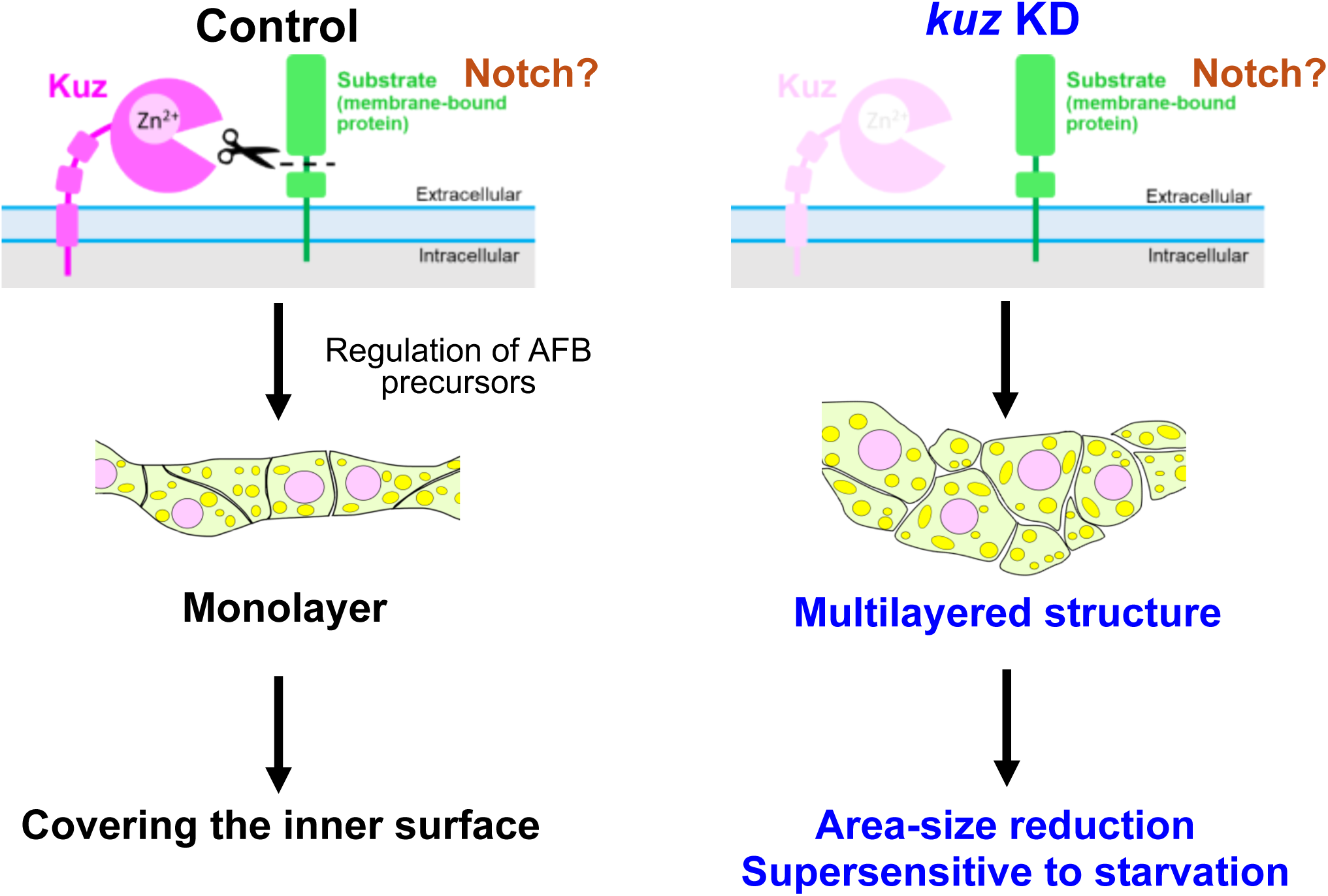
Roles of the ADAM 10 transmembrane metalloprotease, Kuz, in the tissue architecture and function of AFB. Summary of the AFB phenotypes under control (left) or *kuz* knockdown (right) conditions. See text for details.

The *kuz* knockdown AFB was heavily stained with Nile Red at both one day and one week after adult eclosion, similar to the control AFB (Figure 9A), suggesting that lipid storage was not severely affected by the *kuz* knockdown. Next, we examined the AFB morphology in adults using a two-photon microscope. As described in our ultrastructural analysis above, multicell-layered AFB or a clump of AFB cells were found at the dorsal edge of the tergite in the *kuz* knockdown pharate adults. Consistently, near the dorsal midline, the *kuz* knockdown AFB at 0-4 hr after adult eclosion was thicker than the control AFB as indicated by staining with Bodipy (Figure 9B; Movie S1), although it was difficult to judge whether it was multicell-layered or not due to weak phalloidin signals that were expected to label the cell cortex. Collectively, the locally malformed AFB caused by *kuz* knockdown in pharate adults persisted into the adult stage.

### Knocking down *Notch* (*N*) reduced AFB area coverage like the *kuz* knockdown

To understand the role of Kuz in AFB morphogenesis at the molecular level, it is crucial to identify the key substrates of Kuz. A single-pass transmembrane protein, Notch (N), is one of the best-characterized substrates for Kuz or ADAM10 in various contexts in both *Drosophila* and vertebrates (Weber and Saftig, 2012; Rosenbaum and Saftig, 2024). We verified that one of the two RNAi stocks targeting *N* was capable of reducing *N* expression (Figure S8A, B). Using this stock, we examined whether *N* knockdown in AFB development reduced AFB area coverage before or after precursor cell adhesion (Figure S8C-F). The coverage was reduced significantly after adhesion and during monolayer formation, as was also the case with the *kuz* knockdown (Figure 4), implying the possibility that the Kuz-Notch connection also functions in AFB development.

The cleavage of Notch at its extracellular region by ADAM10 or Kuz is required to initiate the N signaling pathway, which leads to the release of the N intracellular domain (NICD), its translocation into the nucleus, and resultant gene expression of N-responsive genes (Weber and Saftig, 2012; Rosenbaum and Saftig, 2024). Generally, the contribution of the Kuz-dependent cleavage of N can be addressed by investigating whether forced NICD expression is capable of rescuing the *kuz* loss-of-function phenotypes. In fact, in a background of a reduced function of *kuz*, the NICD expression ameliorated a migration defect of border cells in the ovary, suggesting a role of the Kuz-N axis in this developmental context (Wang et al., 2006, 2007). Although we made experimental designs to address a possible contribution of the Kuz-N axis to the AFB morphogenesis, unfortunately, we were unable to establish the genetically complex strains essential for the experiments (data not shown).

### *kuz* or *N* knockdown adults were supersensitive to a starvation stress

Finally, we addressed whether *kuz* or *N* knockdown affected AFB function. For this purpose, we examined the sensitivity of adults with the *kuz* or *N* knockdown AFB to starvation stress, given that fat bodies accumulate not only lipid droplets but also glycogen and protein granules and serves as a nutriment reserve upon fasting (Heier and Kühnlein 2018; Chatterjee and Perrimon 2021). We found that the *kuz* or *N* knockdown adults were significantly supersensitive compared to control adults (Figure 9C). Compared to *srp* knockdown adults, which showed almost complete loss of AFB (Tsuyama et al. 2023; Lei et al. 2023), the phenotypes of *kuz* or *N* knockdown were milder. Nonetheless, these data indicate that AFB function is diminished by *kuz* or *Notch* knockdown. Altogether, our results show that *kuz* knockdown flies exhibited defects in both the tissue architecture and function of the AFB.

## Discussion

To elucidate the mechanisms underlying morphogenesis and functional acquisition of any given tissue, the starting point is to identify the newly born tissue in normal development and clarify its tissue architecture. In this respect, the study of adipose tissue has lagged behind that of other tissues. We have focused on the adult fat body (AFB) of *Drosophila melanogaster* as a model system to study the morphogenesis of adipose tissue. In this study, we demonstrate that Kuzbanian (Kuz), a *Drosophila* homolog of the ADAM 10 transmembrane metalloprotease, is required for this critical step of the monolayer formation. Diminished *kuz* function in the precursor cells caused local multilayering or cell clumping. Furthermore, AFBs with such aberrant architectures exhibit diminished function.

In wild-type adults fed a normal diet, the AFB accumulates lipid droplets, swells, and occupies a significant volume within the body cavity. However, the newly formed AFB within the pharate adult, before eclosion, is merely a thin single-cell layer. It is likely for this reason that this nascent AFB was not captured in a previous study that meticulously described the spatial arrangement of organs during metamorphosis using micro-computed tomography (μCT) (Schoborg et al., 2019). By first identifying the AFB in tissue sections of control pharate adults, we were then able to observe the AFB morphology in *kuz* knockdown pharate adults through histological and ultrastructural analysis, revealing the key structural defects caused by *kuz* knockdown.

Among the phenotypes observed in the *kuz* knockdown pharate adult, we first noticed a reduction in the AFB area coverage. It is conceivable that the reduced area coverage is a direct consequence of the AFB, which normally forms a complete monolayer sheet, becoming locally multilayered or forming clumps, preventing it from spreading into a complete sheet. However, we cannot exclude the possibility that *kuz* knockdown in precursor cells adversely affected their proliferation or migration. This would reduce the total number of cells available to participate in monolayer formation, which could also contribute to the reduced area coverage. We attempted to count the number of precursor cells by expressing a nuclear localized fluorescent protein (Red Stinger; Barolo et al., 2004), but unlike with membrane-bound GFPs (mCD8::GFP and CD4:tdGFP; Lee and Luo 1999 and Han et al 2011, respectively), the nuclei of adjacent oenocytes were also labeled, preventing a selective quantification of AFB precursor cell numbers (data not shown). The multilayering or cell clumping phenotype of the *kuz* knockdown was prominent in the dorsal region and this spatial feature could be related to the role of a FGF ligand, which is expressed in the dorsal epidermis, for presumably directing the migration of the precursor cells (Lei et al., 2023).

How does the knockdown of the *kuz* gene in AFB precursor cells lead to the formation of multilayers and cell clumps? This structure of the *kuz* knockdown AFB is reminiscent of malignant cancer cell clones arising in epithelia, or phenomena resembling epithelial-mesenchymal transition (Lu and Kang 2019; Dongre and Weinberg 2019). However, the electron microscopy images show that the multilayered or clumped cells exhibit features of differentiated adipocytes, such as the formation of lipid droplets. Therefore, malignant transformation or a change in adipocyte cell fate is unlikely.

To understand the mechanism by which *kuz* knockdown causes these structural abnormalities, two approaches will be necessary. One is to perform ultrastructural analysis at the stage before AFB precursor cells adhere to each other (earlier than 65 hr APF), to examine potential interactions with surrounding tissues. In this study, our analysis focused on the AFB after it had finished forming a sheet (73 hr APF), and in control pharate adults at that stage, the AFB does not appear to be in physical contact entirely with surrounding tissues. Precursor cells, both during migration and as they cease migration to initiate adhesion, actively extend and retract filopodia. Such cellular dynamics allows us to predict the possibility that other tissues in the close vicinity of the precursor cells provide scaffolds and attractants for the directed migration and spreading, and the candidates tissues include ventral nest histoblasts that form the adult epidermis, lateral muscles in the pleurite, and oenocytes that predominantly reside in the tergites (Tsuyama et al., 2023). A prerequisite this approach is the technical ability to identify migrating precursor cells under an electron microscope.

The second approach is to search for substrate proteins of Kuz that function in the context of AFB formation, and to examine how their molecular functions are regulated by Kuz. We focused on Notch (N), the most well-studied of its known substrates, and showed that *N* gene knockdown reduced AFB area coverage, similar to *kuz* knockdown. It is necessary to perform electron microscopy on *N* knockdown pharate adults to examine whether the AFB becomes multilayered or forms clumps. With respect to cell migration and tissue architecture, other candidates of key substrates of Kuz in the AFB morphogenesis may include a receptor tyrosine kinase, discoidin domain receptor 1 (DDR1), which controls cell migration on collagen (Shitomi et al., 2015) and is conserved in flies, a GPI-linked septate junction protein, Undicht (Petri et al., 2019), and hypothetical cell-cell or cell-substrate adhesion molecules whose ectodomain shedding prevents clumping but facilitates spreading of the precursor cells (Huycke et al., 2024).

*kuz* knockdown flies were not only defective in the AFB tissue architecture, but also supersensitive to starvation stress, indicating a diminished function of AFB. This raises interesting questions: Is the structural alteration of AFB in the *kuz* knockdown fly the cause of the diminished function? In other words, does the single-cell-thick sheet of AFB in the control fly matter for fulfilling its full function? The structural abnormality caused by *kuz* knockdown may have reduced the total surface area of the AFB, leading to a decreased efficiency in releasing stored energy or hormones during starvation. Similarly, it is also plausible that the multilayering of the tissue limits the accessibility of the inner cells (those not facing the hemolymph) to nutrients or hormones, such as adipokinetic hormone, which promotes lipid mobilization and local calcium waves in AFB during starvation (Koyama et al. 2020; Ahmad et al. 2025). Furthermore, since *Drosophila* has an open circulatory system, the flow of hemolymph into the heart (dorsal vessel) might be obstructed, or the circulation efficiency of hormones and metabolites in the hemolymph might be reduced. In other words, the single-cell-thick sheet structure can be considered an optimized tissue architecture that allows all cells to efficiently access the hemolymph. To address the aforementioned questions, it is a prerequisite to make quantitative comparisons of the numbers of adipocytes and the amounts of stored lipids and other metabolites in the control and the *kuz* knockdown AFBs.

Finally, we discuss the significance of this research in comparison to mammalian adipose tissue. *Drosophila* fat bodies have garnered attention as potential models for pathologies like obesity, owing to their similarities of functions and underlying genetic programs with mammalian white adipose tissue (Alfa and Kim 2016; Musselman and Kuhnlein 2018; Chatterjee and Perrimon 2021). In addition, our study has demonstrated that the tissue architecture of AFB is also under the control of a genetic program, that Kuz is one member of this program, and that abnormalities in this architecture, caused by *kuz* knockdown, are accompanied by a decline in tissue function. Just as the *Drosophila* AFB adopts a characteristic monolayer structure, at least during its formation, it is reported in mice that subcutaneous adipose tissue is organized into discrete lobules that are surrounded by connective tissue, suggesting a repetitive structural unit (Chi et al., 2018; Estève et al., 2019). Future studies are warranted to determine what kind of genetic program regulates the architecture of mammalian adipose tissue, whether evolutionarily conserved proteins, including ADAM10/Kuz, play important roles in this program, and furthermore, whether tissue architecture is also critical for physiological function in mammals.

## Experimental Procedures

### Fly strains and husbandry

Flies were raised and maintained at 25°C on standard cornmeal food in plastic vials (Watanabe et al., 2017). Fly strains were obtained from the Bloomington Drosophila Stock Center, KYOTO Stock Center, Vienna Drosophila Resource Center, and published resources. The gene names, fly strains, plasmids, and exact genotypes are summarized in Tables S1 and S2. All relevant information is in FlyBase (Öztürk-Çolak et al., 2024). Precise staging of pharate adults was defined as after puparium formation (APF) at 25°C unless described otherwise. *CD4::tdGFP* (C. Han et al., 2011) and hairpin RNAs were expressed by either *c833-Gal4+ppl-4kb-Gal80+tub-Gal80[ts]* (Tsuyama et al., 2023) or *OK6-Gal4* (Lei et al., 2023) to image the AFB precursor cells and the AFB during metamorphosis and to knock down genes primarily in those cells. *tub-Gal80[ts]* was used for timed control of Gal4 activation (McGuire et al., 2004). The Gal80[ts] protein was a temperature-sensitive mutant of Gal80, which inhibits Gal4 activity under permissive temperatures. After crossing *c833-Gal4+ppl-4kb-Gal80+tub-Gal80[ts]* with the hairpin RNA stock, vials were maintained at a permissive temperature (19°C) to inhibit Gal4 made from *c833-Gal4* in embryonic and larval stages, and prepupae were collected for over two-hour periods (0-2 hr APF at 19°C). These flies were transferred to a restrictive temperature (29°C). In contrast, all steps were conducted at 25°C when *OK6-Gal4* was used. In knocking down *N*, we used *OK6-Gal4* and *elav-Gal80* together to avoid gene knockdown in neurons (Lei et al., 2023). In almost all experiments, the flies were sexed, and females were used for analyses (Table S2).

### Generation of GFP-tagged Kuz stocks

The DNA fragment containing EGFP-FIAsH-StrepII-TEV-3xFlag was amplified using the pBS-KS-attB1-2-PT-SA-SD-1-EGFP-FIAsH-StrepII-TEV-3xFlag plasmid (*Drosophila* Genomics Resource Center Stock 1306) as a template. *GFP-tagged kuz* transgene was created by φC31-mediated genomic cassette exchange with the MiMIC cassette of *MI{MIC}kuzMI9225* strain (Bloomington Drosophila Stock Center 52086; see Figure S2A; Nagarkar-Jaiswal et al. 2015), which was performed by WellGenetics Inc. (Taipei, Taiwan).

### Genome-wide association (GWA) analysis

The rates of AFB developmental defects among 39 representative DGRP lines were previously assessed (Figure S1; Tsuyama et al., 2023). Our GWAS workflow was essentially as described previously (K. Li et al., 2023). Briefly, the phenotypic data were submitted to the DGRP2 web-based analysis pipeline (previously available at http://dgrp2.gnets.ncsu.edu; Huang et al., 2014; MacKay et al., 2012). We defined GWA-significantly associated genetic variants by a false discovery rate of 0.01, which corresponds to a *p*-value of 1.01 × 10^−6^ in our data. Data were visualized using a custom program written in MATLAB (The MathWorks, Inc., Natick, MA, USA).

### Light microscopes

Images of whole adults and dissected abdomens unstained (the far left and second from the left, respectively, in Fig. 1C) were acquired using a digital camera (DP21, Olympus, Japan) attached to a stereo microscope (SZX7, Olympus, Japan). Images of Oil Red O staining and histological sections were acquired using digital cameras (AxioCam color, Zeiss; Digital Sight 1000, Nikon, Japan) attached to an upright microscope (Eclipse E800, Nikon, Japan). For confocal imaging, flies were imaged using a Nikon-C1 confocal laser scanning microscope equipped with an inverted microscope (Eclipse Ti, Nikon, Japan) or an upright microscope (Eclipse E800, Nikon, Japan). Confocal imaging of whole-mount live female pharate adults was performed as previously described (Tsuyama et al., 2023). Two-photon imaging was performed using an Olympus FV1000MPE-IX83 multiphoton laser scanning microscope, and the acquired images were processed using Imaris x64bit 10.0.0 (OXFORD Instruments, UK).

### Histological analysis

Pharate adults were fixed in Bouin’s solution (HT10132, Sigma-Aldrich, Germany) for 4 hr and rinsed with 70% ethanol. The samples were dehydrated using a graded ethanol series, cleared in chloroform, and embedded in paraffin. Serial frontal sections (3–4 µm thick) were cut at 50 µm intervals and stained with hematoxylin (1.15938.0025, Merck, Germany) and eosin Y (1.15935.0025, Merck, Germany) (H&E) using standard methods.

### Ultrastructural analysis

Pharate female adults were permeabilized with acetone for 5 s and fixed in 5% glutaraldehyde (17003-92, Nacalai Tesq, Japan) overnight (Butterworth & Forrest, 1984). Resin embedding and serial sectioning were performed as previously described (Koyanagi et al., 2024). Briefly, after washing with 0.1 M Phosphate buffer (PB), the samples were post-fixed with 2% (w/v) osmium tetroxide at room temperature for 1.5 hr. Samples were then dehydrated in a graded series of ethanol followed by 100% propylene oxide and embedded in epoxy resin (Luveak-812, Nacalai Tesq, Japan) according to a standard procedure. Serial sections (250 nm thickness) were cut with a diamond knife (SYM NV3045 Ultra, SYNTEK, Japan) using an ultramicrotome (ARTOS 3D, Leica, Germany). The sections were collected on a clean silicon wafer strip held by a micromanipulator (MN-153, NARISHIGE, Japan). The sections were stained with 2% (w/v) aqueous uranyl acetate for 20 min and Reynolds’ lead citrate for 2 min. Backscattered electron (BSE) images of the serial sections were acquired using field emission scanning electron microscopy (FE-SEM; JSM-7900F; JEOL, Japan).

### Immunohistochemistry

For immunostaining AFB in pharate adults or adults, the abdomens were dissected and fixed as described previously (Tsuyama et al., 2023). Fixed tissues were permeabilized with 3% Triton X-100 in PBS for 15 min and washed in PBS five times. After blocking with PBST (PBS containing 0.3% Triton X-100) plus 2% bovine serum albumin (BSA) for 60 min, specimens were incubated with chicken anti-GFP antibody (1:300; ab13970, Abcam, UK) overnight at 4 °C. After five washes in PBS, the samples were incubated with Alexa 488-conjugated donkey anti-chicken IgY antibody (1:500; 703-545-155, Jackson ImmunoResearch Labs, USA) for 30 min at room temperature. The samples were mounted in 50% glycerol in PBS with a piece of plastic tape as a spacer to avoid crushing the specimens.

To identify developing AFB in histological sections, the sections of the pharate adult that expressed GFP in the AFB were processed as follows: after deparaffinization, endogenous peroxidase activity was blocked using 0.3% H_2_O_2_ in methyl alcohol for 30 min. The glass slides were washed in PBS six times and mounted with 1% normal serum in PBS for 30 min. Subsequently, chicken anti-GFP antibody (1:1200; ab13970, Abcam, UK) was applied overnight at 4 °C. The slides were then incubated with biotinylated goat anti-chicken IgY antibody (H+L) (1:300; BA-9010-1.5; Vector Laboratories, USA) for 40 min, followed by washing with PBS six times. Avidin-biotin-peroxidase complex (ABC-Elite, Vector Laboratories, USA) was diluted 1:100 in BSA-containing PBS and applied for 50 min. After washing in PBS six times, the coloring reaction was carried out with 3,3’-diaminobenzidine tetrahydrochloride (DAB), and the nuclei were counterstained with hematoxylin (Toda et al., 1999).

For immunostaining of the central nervous system (CNS), wandering third-instar larvae were dissected in PBS and fixed with 3.7% formaldehyde + 0.05% Triton-X100 in PBS overnight at 4 °C. After four washes with PBST, the samples were blocked with 2% BSA in PBST for 30 min at room temperature. The samples were then incubated with chicken anti-GFP antibody (1:1000; ab13970, Abcam, UK) for 3 d at 4 °C. After five washes with PBST, the samples were incubated with Alexa 488-conjugated donkey anti-chicken IgY antibody (1:1000; 703-545-155, Jackson ImmunoResearch Labs, USA) for 2 h at room temperature. After further washing, the samples were mounted using ProLong Glass Antifade Mountant (Thermo Fisher, USA). For immunostaining of wing discs, wandering third-instar larvae were dissected and fixed in the formaldehyde solution described above for 15 min at room temperature. The samples were washed, blocked, and incubated with mouse anti-Notch intracellular domain antibody (1:200; C17.9C6, Developmental Studies Hybridoma Bank, USA) overnight at 4 °C. The secondary antibody was goat anti-mouse IgG antibody (A21424, Molecular Probes, USA).

### Staining lipid droplets and cell cortex

The abdomens of pharate adults and adults were dissected and fixed as in the immunohistochemistry and stained with Nile Red (1 μg/ml; 144-0811, WAKO, Japan) for 3 min or Oil Red O (0.18%; O0625, Sigma-Aldrich, Germany) for 20 min at room temperature. For two-photon imaging, female adults at 0-4 hr after eclosion were dissected and stained with BODIPY 493/503 (25 ng/ml: D3922, Invitrogen, USA) for 15 min and Phalloidin 647 (1x solution; A22287, Thermo Fisher, USA) for 2 h at room temperature.

### Western blotting

CNS was dissected out of 10 wandering 3rd instar larvae, placed in 15 μl 1x SDS sample solution containing 5% 2-Mercaptoethanol, sonicated using Branson Sonifier 450, and boiled for 4 min. The solubilized proteins were separated by SDS-PAGE (5-20% gradient; 27GM, FUJIFILM Wako Pure Chemical Corporation, Japan) and wet transferred to polyvinylidene difluoride (PVDF; ISEQ00010, MERCK, Germany). The amount of protein per lane was equivalent to that from the 10 tissues. The membrane was probed with rabbit anti-GFP antibody (1:2000; A11122, Invitrogen, USA) or rat anti-Dα−catenin DCAT1 (Oda et al., 1993). The secondary antibodies used were horseradish peroxidase (HRP)-conjugated donkey anti-rabbit IgG antibody (1:2000; NA934-1ML, Cytiva, USA) and HRP-conjugated goat anti-rat IgG antibody (1:2000, ab205720, Abcam, UK). A chemiluminescence assay kit (Chemi-Lumi One Super; 02230-30, Nacalai Tesq, Japan) was used, and signals were detected using ImageQuant LAS 4000 (Cytiva, USA).

### Quantification of AFB area coverage

The area of AFB tissues in fluorescence images was measured using the Weka Trainable Segmentation (Arganda-Carreras et al., 2017) plugin in ImageJ, as described previously (Tsuyama et al., 2023). The AFB area coverage was calculated as the percentage of the area size of the tergite AFB to the whole tergite area size.

### Starvation assays

0-12 hr-old virgin female adults were collected and were reared on standard cornmeal food for 60 h. Subsequently, the female adults were transferred into vials containing 2% agar medium, and numbers of surviving, dead, and censored flies were counted at 12 h intervals. Each condition consisted of 11-13 vials containing 7-13 adults per vial (total 114-126 adults per knockdown of one gene). Survivorship was analyzed using OASIS2 (S. K. Han et al., 2016; https://sbi.postech.ac.kr/oasis2/)

### Statistical analysis

Statistical analyses were performed using the R software. The R codes used in this study are available on request. Statistical significance was set at p < 0.05. The Wilcoxon-Mann-Whitney test was used for two-group comparisons, and Steel’s test was used for multiple comparisons. The log-rank test was used to compare the survival curves and test for statistical significance. The boxes in the boxplots represent the upper and lower quartiles, and the central lines indicate the median. The whiskers extend to the most extreme data points, which are no more than 1.5 times the interquartile range.

## Acknowledgements

We thank K. Takakura and K. Aoki of the Kyoto University Live Imaging Center and iSAL of Kyoto University for their assistance with two-photon imaging. Histopathological procedures were carried out at the Division of Histological Study, and the electron microscopic study was performed in the Division of Electron Microscopic Study, Center for Anatomical Studies, Graduate School of Medicine, Kyoto University. We also thank A. Nose, M. Enomoto, and J. H. Simpson for providing us with fly strains; R. Tajiri, M. Furuse, Y. Izumi, S. Yonemura, Atsuko Sehara-Fujisawa, Y. Yoshinari, T. Nishimura, T. Koide, K. Saito, T. Maeno, A. Kimura, and S. Nishikawa for technical advice and/or discussion; J. Hejna for polishing the manuscript; M. Futamata, K. Oki, H. Imai, and Y. Niitani for technical and secretarial assistance; the other members of the Uemura laboratory for technical advice and discussion; and H. Matsumoto for the illustrations of pharate adults. The reagents, genomic datasets, and/or facilities were provided by the Bloomington *Drosophila* Stock Center, the Vienna Drosophila Resource Center (VDRC, www.vdrc.at), the TRiP at Harvard Medical School (Grant: National Institutes of Health, Office of the Director R24 OD030002), the *Drosophila* Genetic Resource Center at Kyoto Institute of Technology, and FlyBase (Larkin et al., 2021; Gramates et al., 2022). This work was supported by the Japan Society for the Promotion of Science (JSPS; 23K27179 to T.U.), the Japan Agency for Medical Research and Development (AMED-CREST; JP18gm1110001 to T.U.), K. Kato, and JSPS KAKENHI Grant Number JP22H04926 (24A-012-I22 to T.U.).

---

**FIGURE S1.**
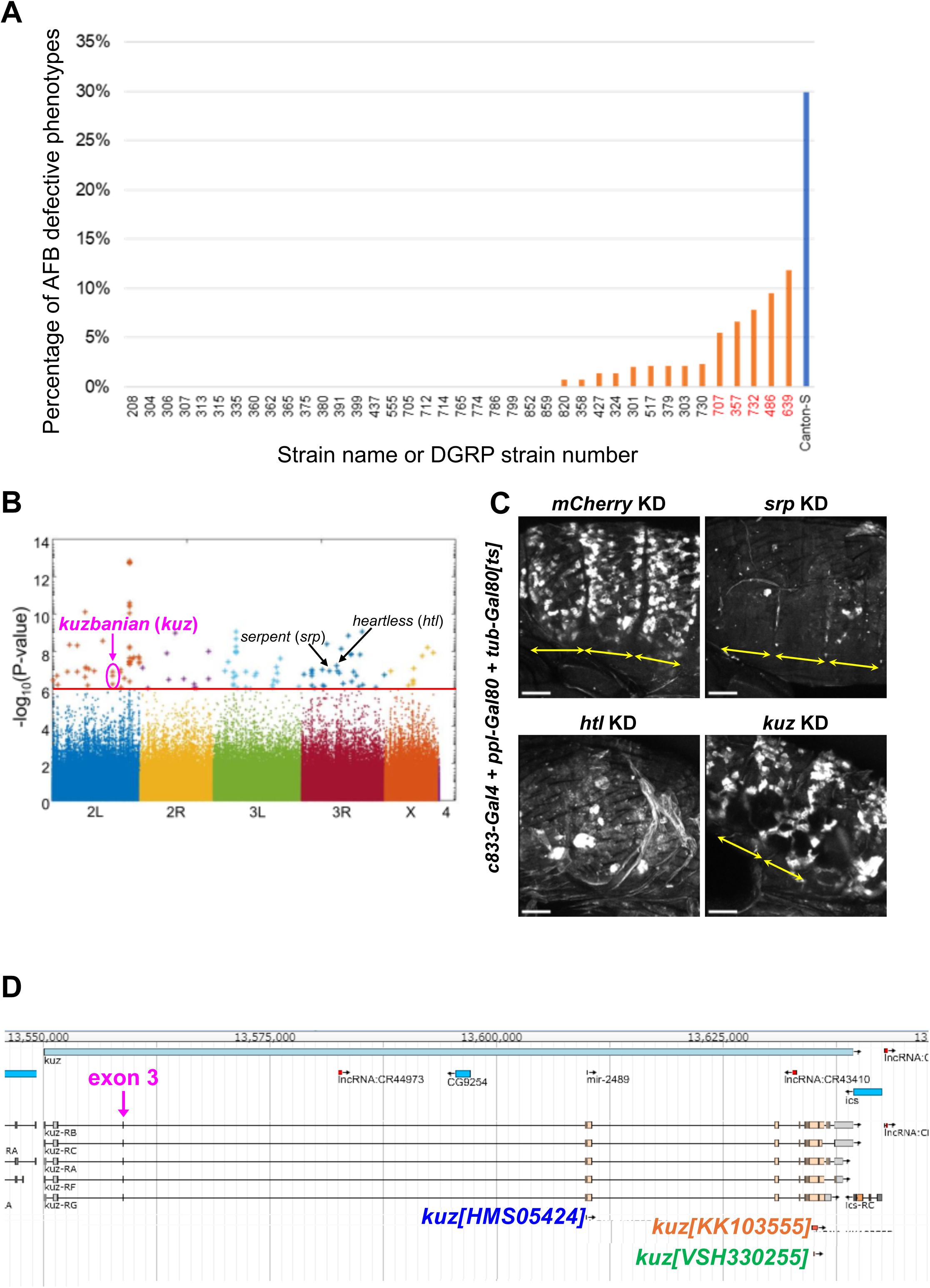
Genome-wide association (GWA) analysis and knockdown screening for genes regulating AFB development. (A) Rates of the AFB-defective phenotype of 39 representative DGRP lines and *Canton-S* in ascending order (n=131 to 158 adults/line, except for n=37 of DGRP-714; Tsuyama et al. 2023). (B) GWA analysis of the AFB defect. The p*-*values(−log10 transformed) are shown on the y-axis. The red line marks the nominal p*-*value threshold (1.01×10–6). Each data point corresponds to an individual genetic variant (single nucleotide polymorphism, deletion, or insertion). Data points are arranged by relative chromosome (genomic) position, and the color code indicates the respective chromosome to which they belong. Genetic variants of *kuzbanian* (*kuz*)*, serpent* (*srp*), and *heartless* (*htl*) genes are indicated by the arrows and circles. (C) Representative results of knocking down candidate genes in AFB precursors. Knocking down and cell labeling were performed using *c833-Gal4+ppl-4kb-Gal80+tub-Gal80[ts]* in combination with *CD4::tdGFP*. The control short hairpin RNA for knockdown (KD) was *mCherry*. Flies were aged at 19°C to inhibit Gal4 expression and shifted to 29°C (see details in “Fly strains and husbandry” in Experimental Procedures). Images are whole-mount lateral views of abdomens at 73hr APF at 29°C, which was equivalent to a stage shortly before eclosion. The yellow arrows indicate the individual abdominal segments. Regarding the patchy *CD4::tdGFP* signal in the control AFB, see the legend of Figure 3B. n=3 for each group. Scale bars: 100 µm. (D) The genomic structure of the *kuz* gene and target sequence regions for the knockdown. Total 6 genetic variants are significantly associated with the defective AFB development, and all of them are located in the 2.06 kb genomic region between 2L_13558032 and 2L_13560082, which includes the third exon. Exact nucleotide positions and sequences of individual variants are shown in Table S3. The target sequence of *kuz[KK103555]* (677 bp) includes that of *kuz[VSH330255]* (21 bp). Stocks were indicated on captured images from the JBrowse viewer at FlyBase (Öztürk-Çolak et al., 2024).

**FIGURE S2.**
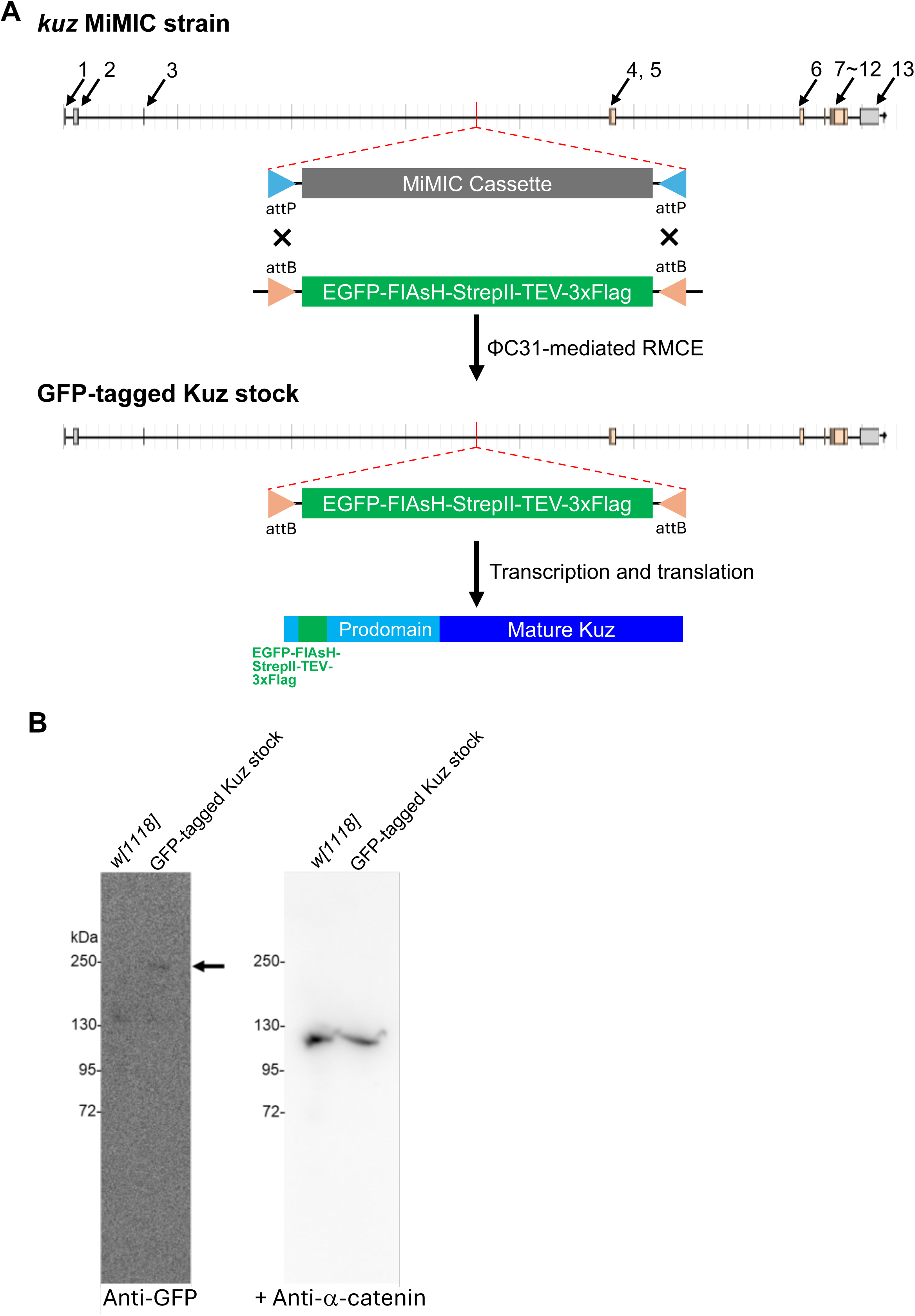
Generation of a “MiMIC GFP-tagged Kuz” stock and detection of the GFP-tagged protein. (A) Schematic representation of the generation of a *GFP*-tagged *kuz* transgene. The genomic structure of the *kuz* locus in *MI{MIC}kuzMI9225* strain (Bloomington Drosophila Stock Center 52086) is shown at the top. The numbers represent exons, and the MiMIC cassette is inserted into the longest intron by ϕC31 recombinase-mediated cassette exchange (RMCE). The resultant protein is tagged with EGFP and Flag, presumably in its prodomain, and the molecular weight of the entire immature protein is predicted to be between 199 and 214 kDa, depending on the tagged isoforms A, B, C, G, and F (see Figure S1D). (B) Western blot analysis of larval central nervous system (CNS) lysates of *w[1118]* and the MiMIC GFP-tagged Kuz stock. We prepared the lysates of larval CNS because it highly expresses *kuz* (Brown et al. 2014). The blot was probed with chicken anti-GFP antibody and subsequently with goat anti-chicken IgY antibody conjugated with horseradish peroxidase (HRP). After the signal detection, the blot was reprobed with rat anti-Dα-catenin antibody (a loading control), without prior stripping the anti-GFP antibody or inactivating HRP using hydrogen peroxide water (as indicated by “+” in front of “Anti-α-catenin”), and then incubated with goat anti-Rat IgG antibody conjugated with HRP. A faint signal around or slightly smaller than the 250 kDa marker (arrow) could be that of the MiMIC GFP-tagged Kuz precursor protein. Under a confocal microscope, GFP signals were detected in the cortex glia (compare “*lexA* KD in GFP-tagged Kuz stock” with “*w[1118]*” in Figure 3E). However, GFP signals were hardly seen in the AFB development (data not shown), presumably because of low expression below the level of detection. We also tested a previously reported “GFP-tagged Kuz in BAC” stock (Dornier et al. 2012). This stock contains a Kuz-GFP transgene, where GFP is tagged at the carboxyl terminus of the Kuz isoforms A and B. A single copy of this transgene rescues the lethality of *kuz* mutant flies, indicating that the Kuz-GFP proteins are functional (Dornier et al. 2012). We found GFP signals in the AFB neither by live imaging of whole-mount pharate adults nor by immunohistochemistry of dissected and fixed abdomens (data not shown). This may be due to a very low amount of Kuz-GFP in AFB, as assumed by the low GFP signals in this stock even in larval imaginal tissues and central nervous system where *kuz* transcript levels are moderate and moderately high, respectively (Dornier et al. 2012; Brown et al. 2014).

**FIGURE S3.**
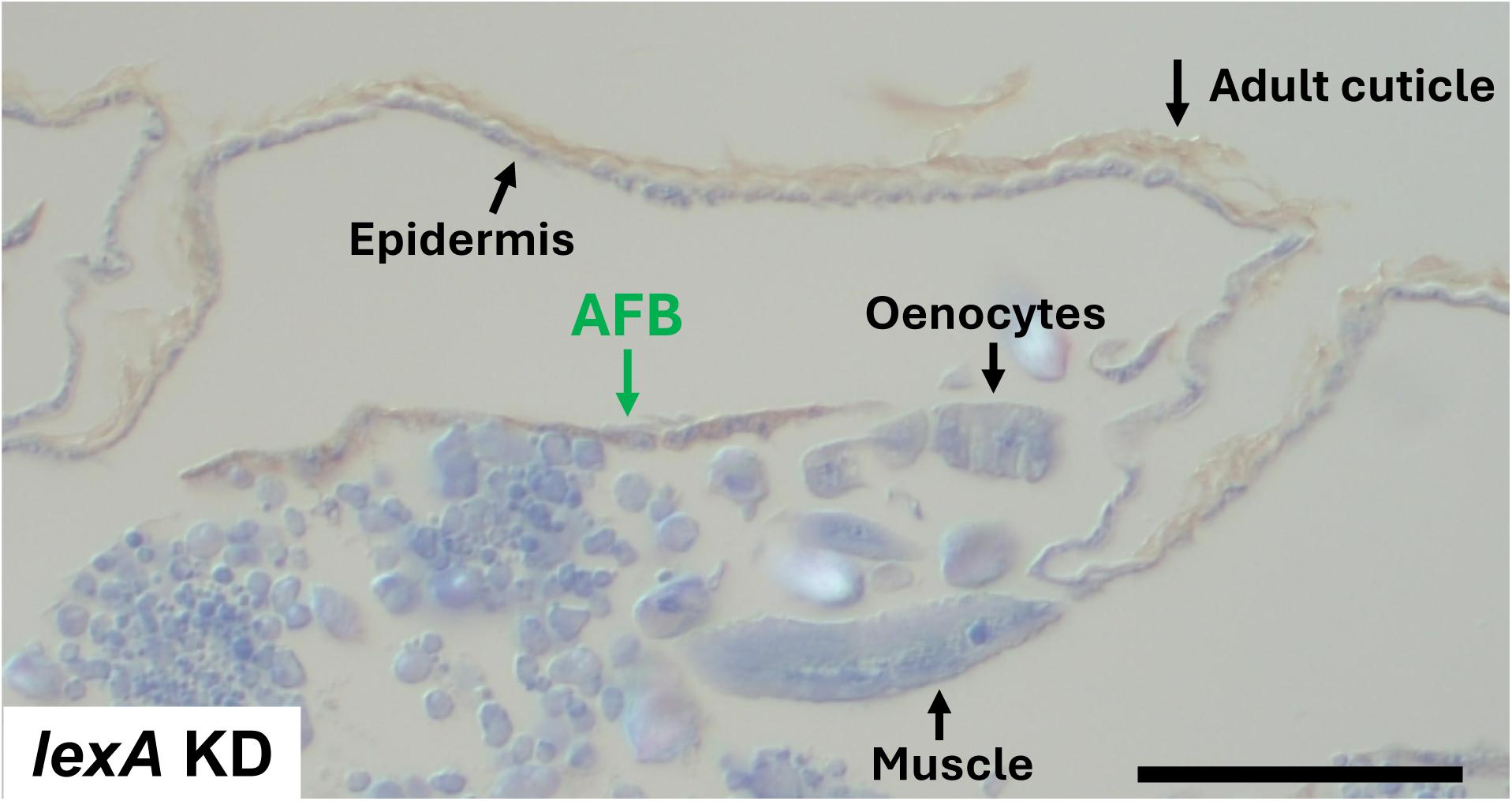
Immunodetection of GFP-expressing AFB on a frontal section at the dorsal level. A frontal section of a 73 hr-APF pharate adult with a control genotype (*lexA* KD) and expressing *CD4::tdGFP* in AFB. This section is located similarly to that shown in Figure 5C. Immunohistochemistry using an anti-GFP antibody colored the thin-layer tissue (AFB) brown, whereas the epidermis, muscle, and oenocytes were blue due to staining with hematoxylin. The adult cuticle was brown in a non-specific manner. n=8.

**FIGURE S4.**
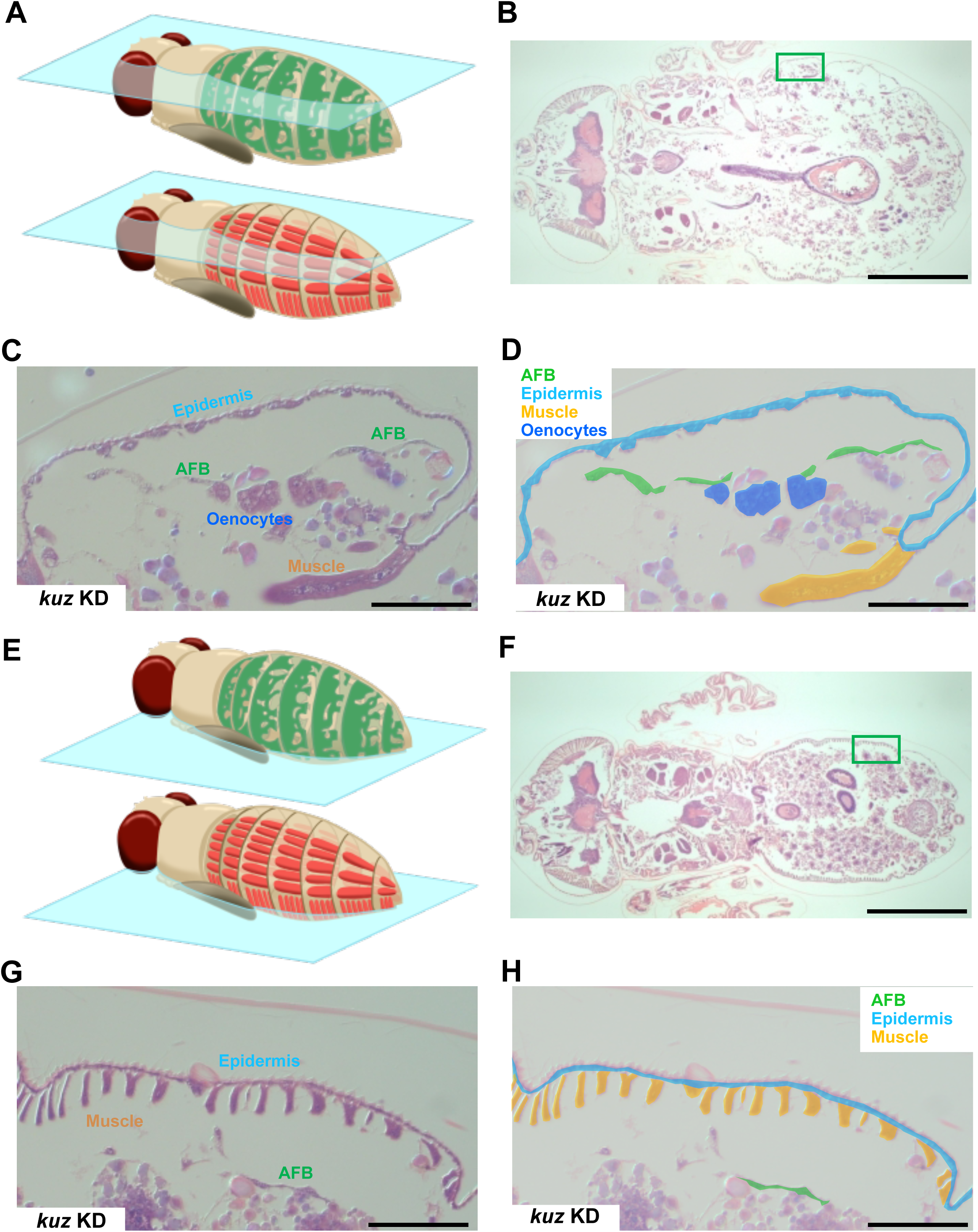
Histological analysis of the *kuz* knockdown pharate adults. (A, E) Schematic illustrations of AFB (green) and muscles (orange) in the abdomens of pharate adults and frontal sections at the dorsal and ventral level (semitransparent cyan boards in “A” and “E,” respectively). (B-D) Images at the dorsal level that were taken from a pharate adult different from the one in Figure 5G and 5H. (F-H) Images at the ventral level that were taken from a pharate adult different from the one in Figure 6G and H. The body surface areas of the abdomens (green boxes in B and F) are enlarged, and the identified tissues are labeled (C, G) or pseudocolored to indicate individual tissues (D, H) as in Figure 5 and 6. Scale bars: 500 µm in B and F; 50 µm in C-D and G-H.

**FIGURE S5.**
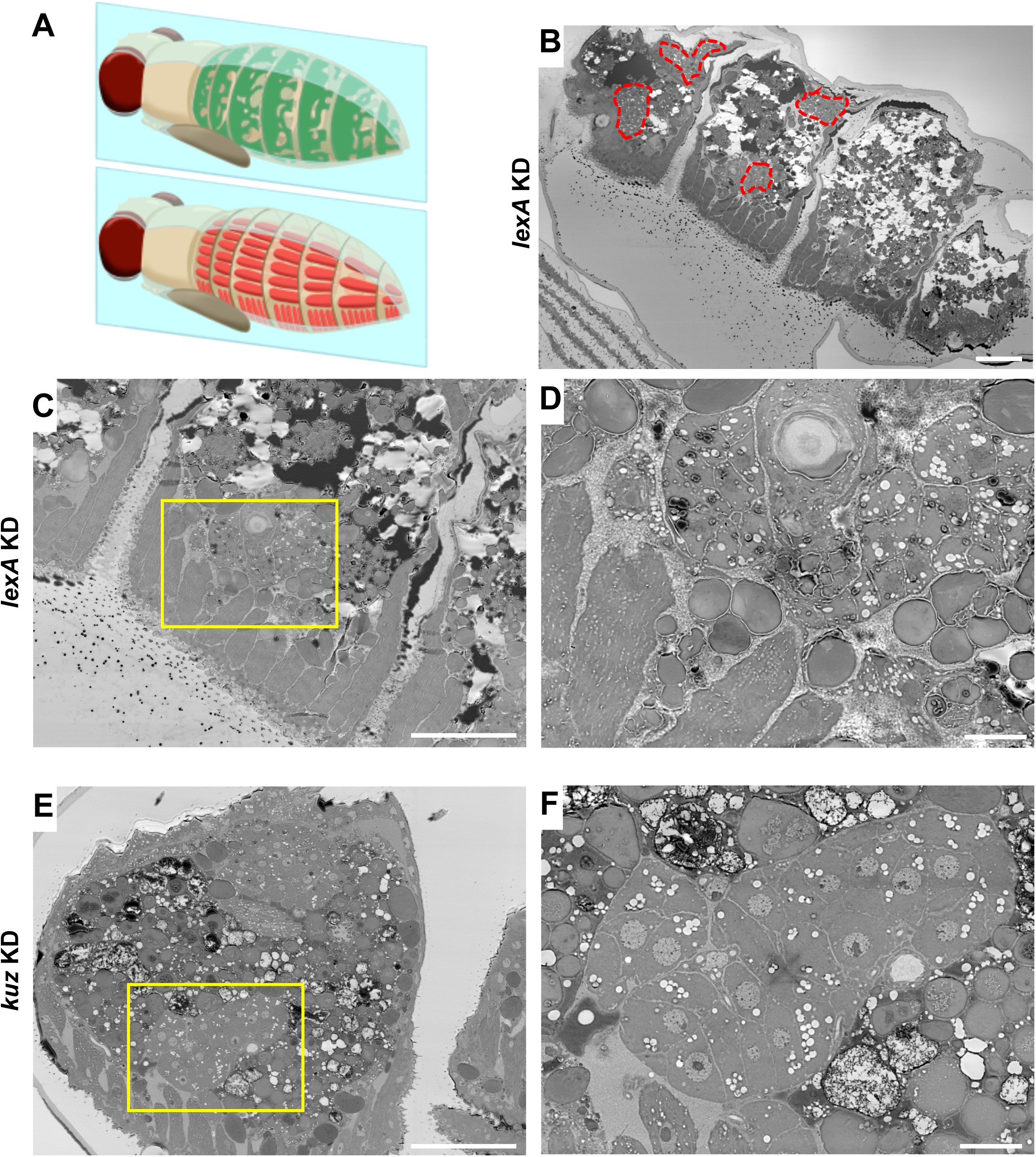
Ultrastructural analysis of AFB on sagittal sections of the control or the *kuz* knockdown pharate adults. (A) Schematic illustrations of AFB (green) and muscles (orange) in the abdomens of pharate adults and sagittal sections (semitransparent cyan boards). (B-F) Representative BSE images of sagittal sections in the lateral-to-ventral region of the abdomen (“Pleurite” in Figure 2C) at 73 hr APF. (B-D) Images of the control pharate adult. (B) AFB was found in areas surrounded by red dashed lines. (C, D) AFB in “B” is zoomed in at two distinct magnifications and the yellow box in “C” is zoomed in “D.” (E, F) Images taken from a *kuz* knockdown pharate adult. The yellow box in “E” is zoomed in “F.” The magnification of “E” is equal to that of “C,” and the ratio of “F” is equal to that of “D.” In all images, anterior is to the left. In the lower-left corners of “C”-“F”, muscle bundles running along the dorsal-ventral axis are seen. Scale bars: 50 µm in (B), (C), and (E); 10 µm in (D), (F). n (microscope fields)=3 (*lexA* KD) and 4 (*kuz* KD). n (flies)=2 for each genotype.

**FIGURE S6.**
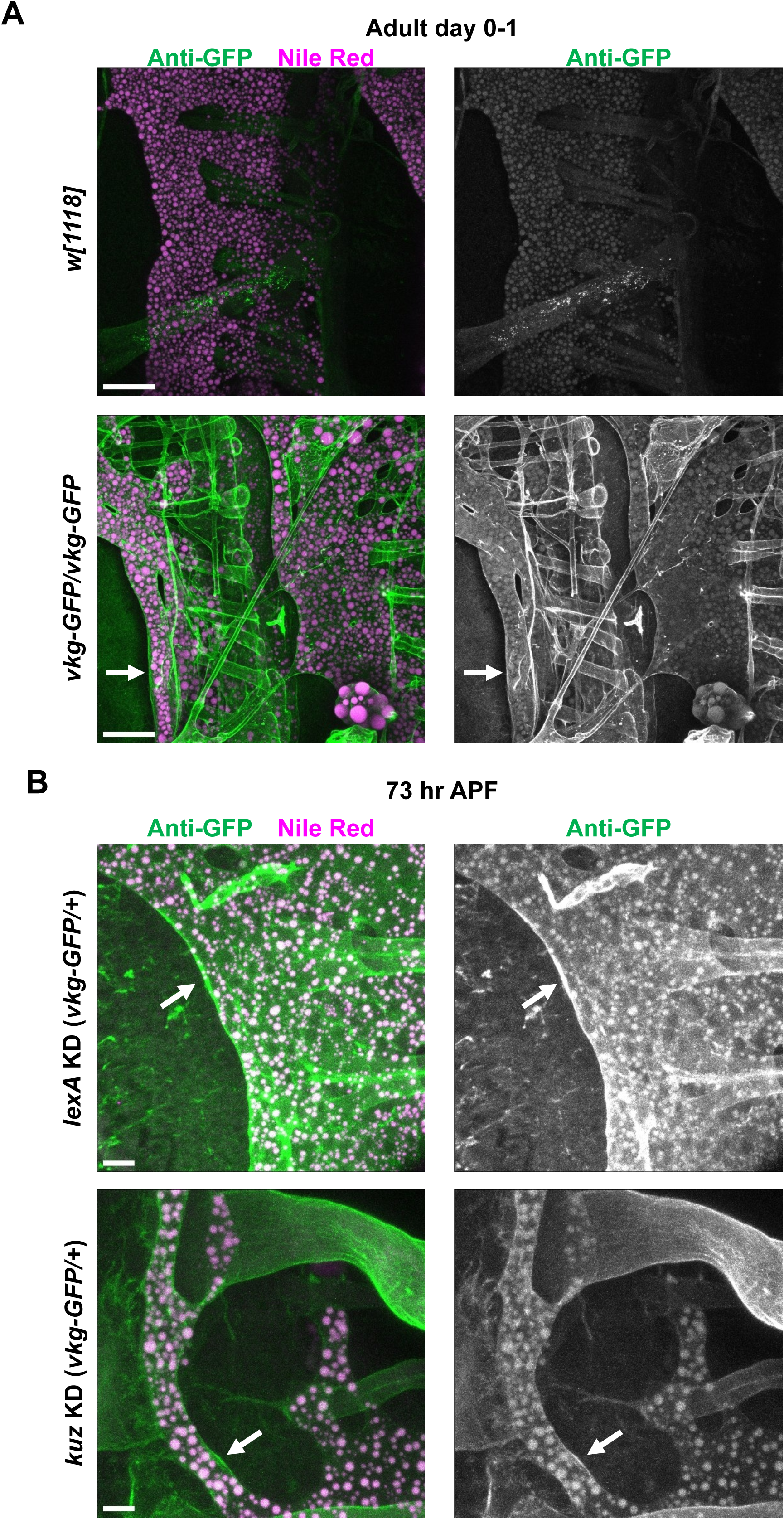
Signals of collagen IV in AFB. Representative images of dissected abdomens stained for GFP (green) and Nile Red (magenta or white). (A) Adults (day 0-1). An adult that did not express GFP in any of the tissues (*w[1118]*; n=2) and an adult homozygous for collagen IV GFP-trap, *viking[G454]* (*vkg-GFP*/*vkg-GFP*; n=3). (B) Pharate adults (73 hr APF). Arrows indicate Vkg-GFP signals on the tissue surface of the AFB. All images were taken near the dorsal midline, where the AFB coverage was severely impaired in the *kuz* knockdown pharate adult (see Figure 2A). The anterior is to the left and the dorsal is upwards. In the larval fat body (LFB), collagen IV is concentrated in the intercellular space, which contributes to inter-adipocyte adhesion and continuity of the LFB tissue layer (Dai et al. 2017). We detected the GFP signal in AFB, which enclosed the AFB layer (arrows in “A” and “B”) and may be associated with tracheal branches or muscles, but did not find obvious signals in the intercellular space within the limits of our observation in either genotype. In the GFP channel images both in (A) and (B), fluorescence signals were also detected in lipid droplets, due to bleed-through signals emitted from NileRed. n=2 (*lexA* KD) and 3 (*kuz* KD). Scale bars: 50 µm in (A); 100 µm in (B).

**FIGURE S7.**
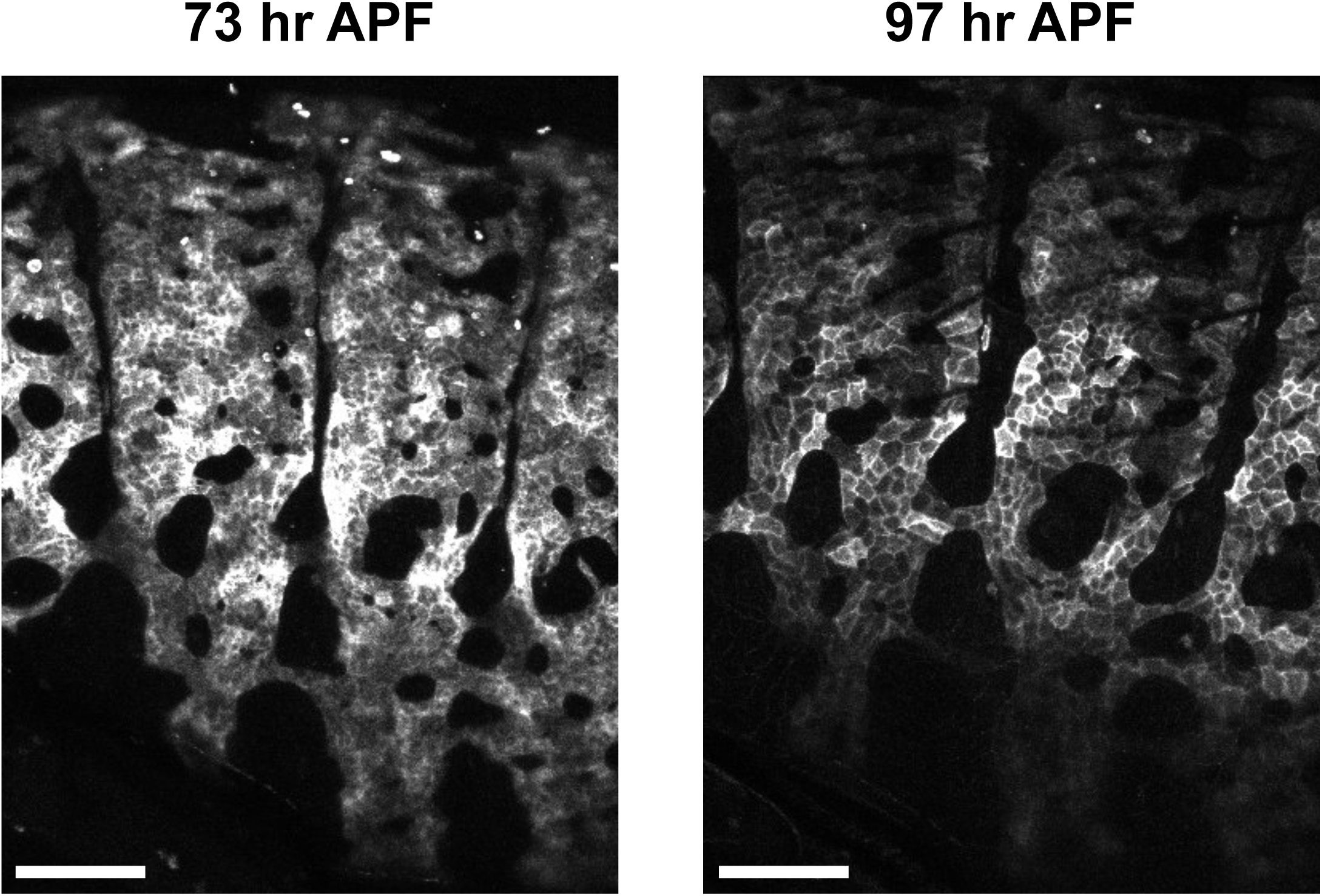
Expression of *OK6-Gal4* at late pharate adult stages. Lateral views of the abdomen of the same whole-mount pharate adult at two distinct timepoints between AFB monolayer formation and eclosion. *OK6-Gal4*-dependent gene expression was visualized by using *UAS-CD4::tdGFP.* Images were acquired and processed under the same conditions. GFP signals became weaker at 97 hr APF, especially in intracellular (non–plasma-membrane) regions, suggesting a decrease in Gal4 activity late in metamorphosis. Scalebars: 100 µm. n = 3.

**FIGURE S8.**
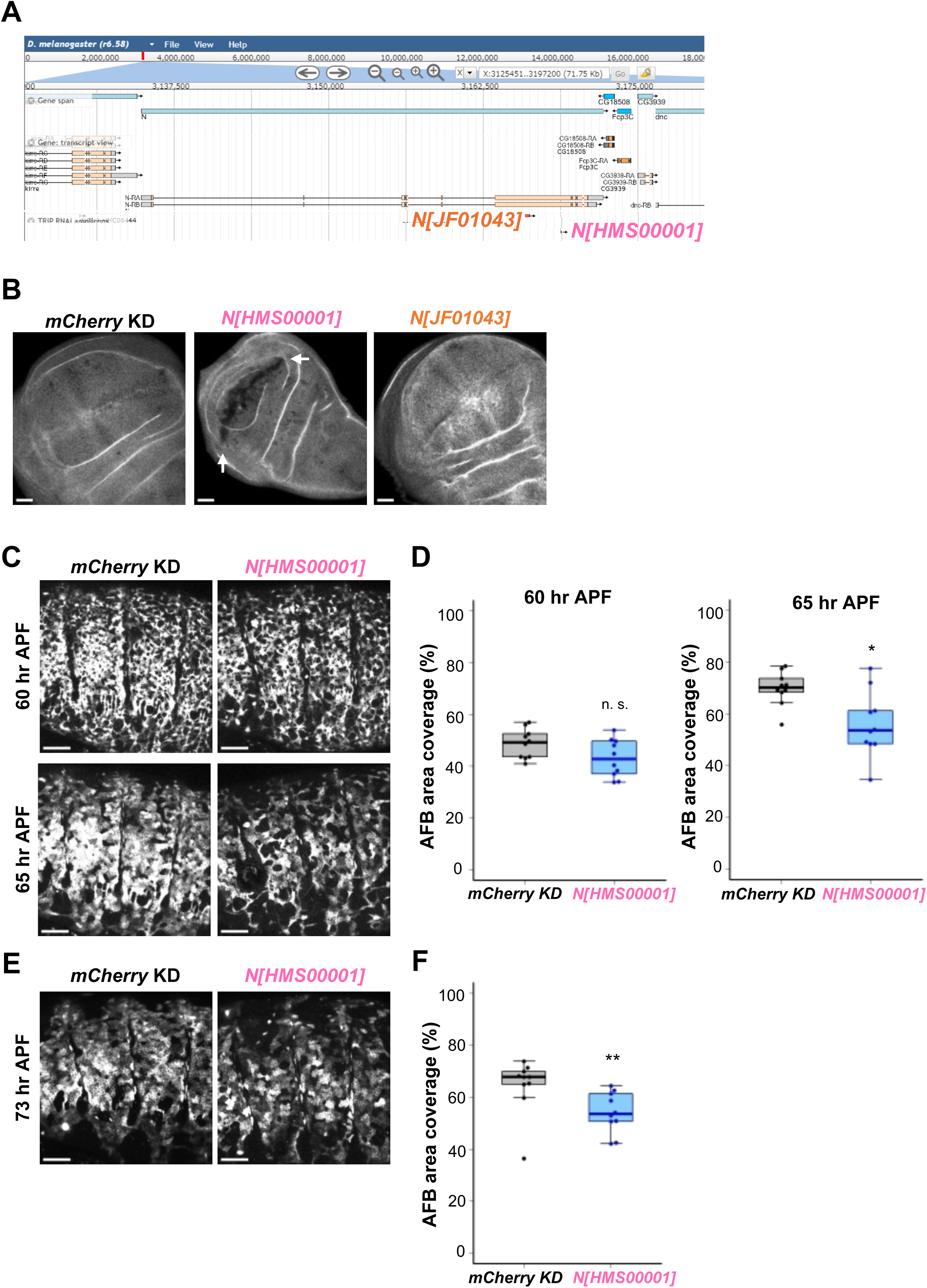
Reduction in the AFB area coverage in pharate adults when *Notch* (*N*) was knocked down. (A) Regions of target sequences for knocking down *Notch (N)* in the fly genome. Stocks are indicated in the captured images from the JBrowse viewer at FlyBase. (B) Immunostaining of wing imaginal discs using an anti-Notch antibody. To knock down *N* along the presumptive wing margin, two stocks were employed with *c96-Gal4* and only one of them, *N[HMS0001]*, was found to be effective (dark margin marked with arrows). The control short hairpin RNA was *mCherry*. n=6 (*mCherry* KD), 4 (*N[HMS0001]* KD), and 5 (*N[JF01043]* KD). (C-F) (C and E) Lateral views of the abdomens of whole-mount pharate adults of the control (*mCherry* KD) and *N* knockdown (*N[HMS0001]*) during (C) and after (E) precursor cell adhesion and monolayer formation. In all images, the anterior is to the left and dorsal is upwards. (D and F) Quantification of AFB area coverage at individual time points. *p<0.05 and **p<0.01. Wilcoxon-Mann-Whitney test. n=10 for each time point of each genotype. Scale bars: 100 µm.

**Movie S1.**
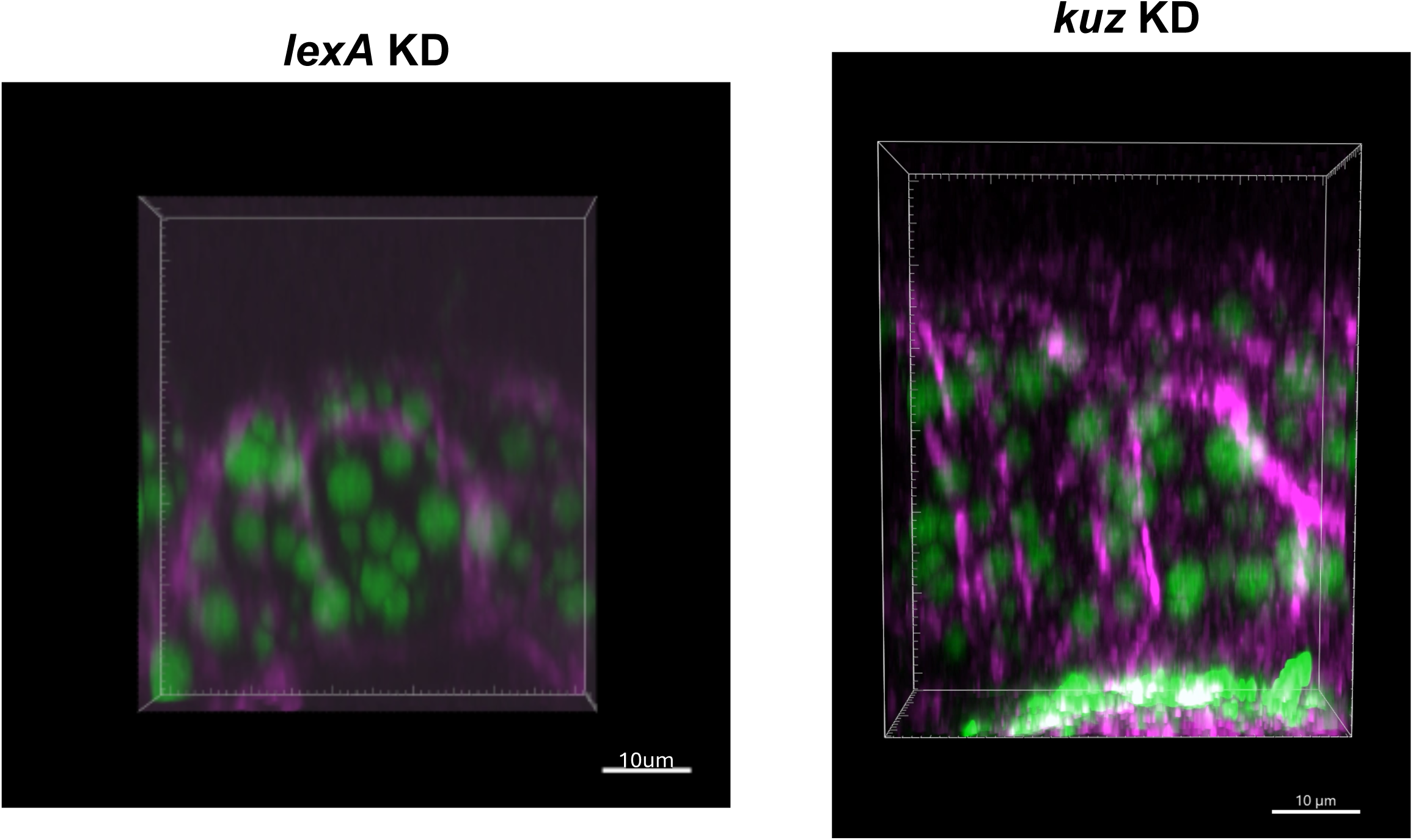
AFB near the dorsal midline. Abdomens of control or *kuz* knockdown adults (0-4 hr after eclosion) were dissected, stained with BODIPY (green) and phalloidin 647 (magenta), and imaged as described in Figure 9B. Dorsal is to the left, anterior is to the front, and the epidermis is downwards in the first frame. Scale bars: 10 µm.

**Table S1.** Genes that were knocked down in AFB precursors and the RNAi lines employed.

| Gene<br>Ontology | Target<br>gene | BDSC<br>strain No. |
| --- | --- | --- |
| Cell<br>migration | *htl* | 35024 |
|  |  | 57313 |
|  | *sr* | 27701 |
|  |  | 60024 |
|  | *numb* | 31182 |
|  |  | 35045 |
|  | *kuz* | 66958 |
|  | *haf* | 28528 |
|  |  | 51438 |
|  | *mirr* | 31907 |
|  |  | 42960 |
|  | *Imp* | 34977 |
|  |  | 38219 |
| Cell<br>adhesion | *ago* | 31501 |
|  |  | 34802 |
| Cell<br>migration<br>and cell<br>adhesion | *caps* | 28020 |
|  | *Dscam2* | 51839 |
|  | *CASK* | 27556 |
|  |  | 32857 |
|  | *Gli* | 31869 |
|  |  | 38284 |
| Stem cell<br>regulation | *srp* | 34623 |
|  | *Nedd8* | 33881 |
|  | *CG8273* | 34805 |
|  |  | 60139 |

**Table S2.**
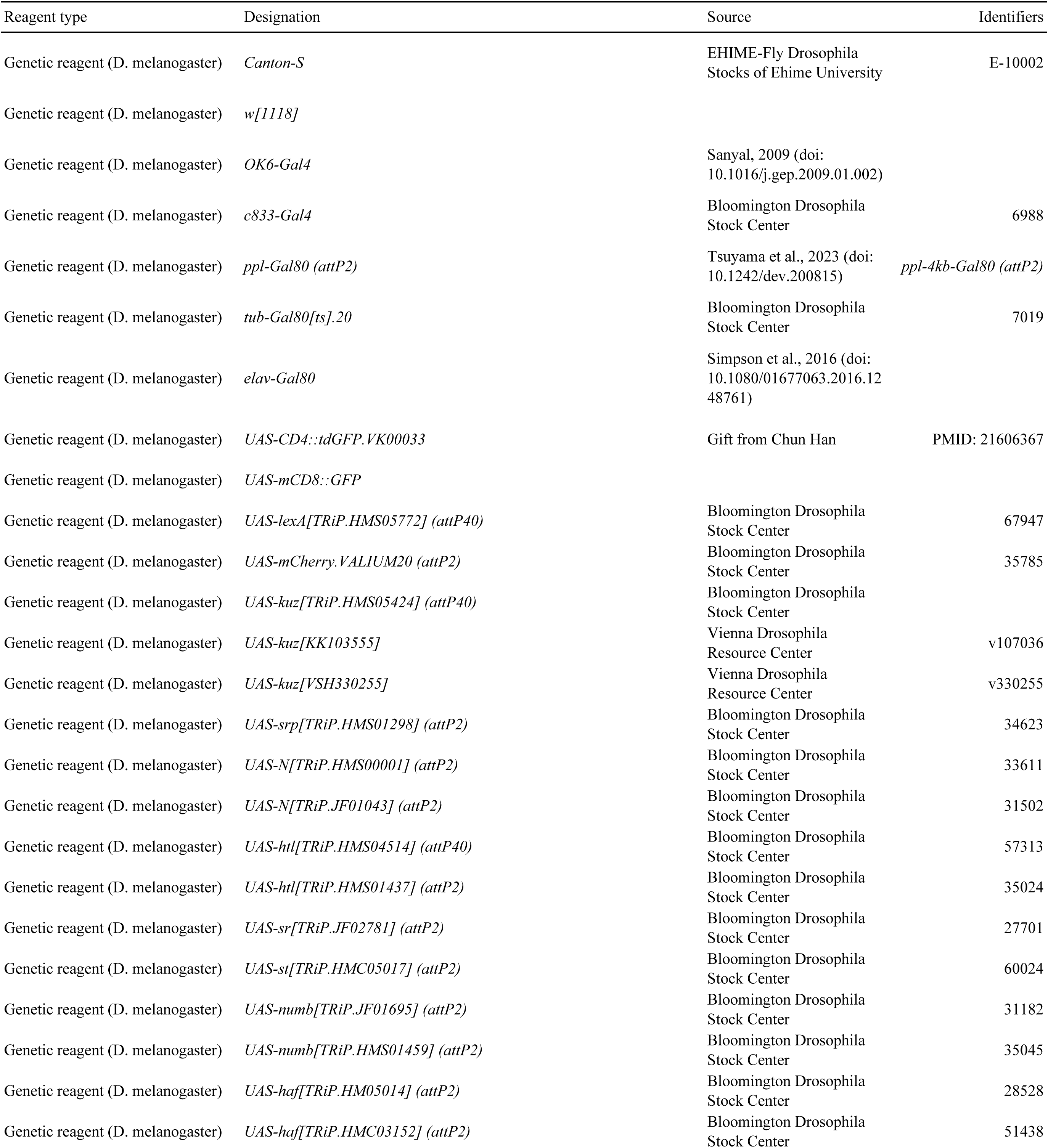
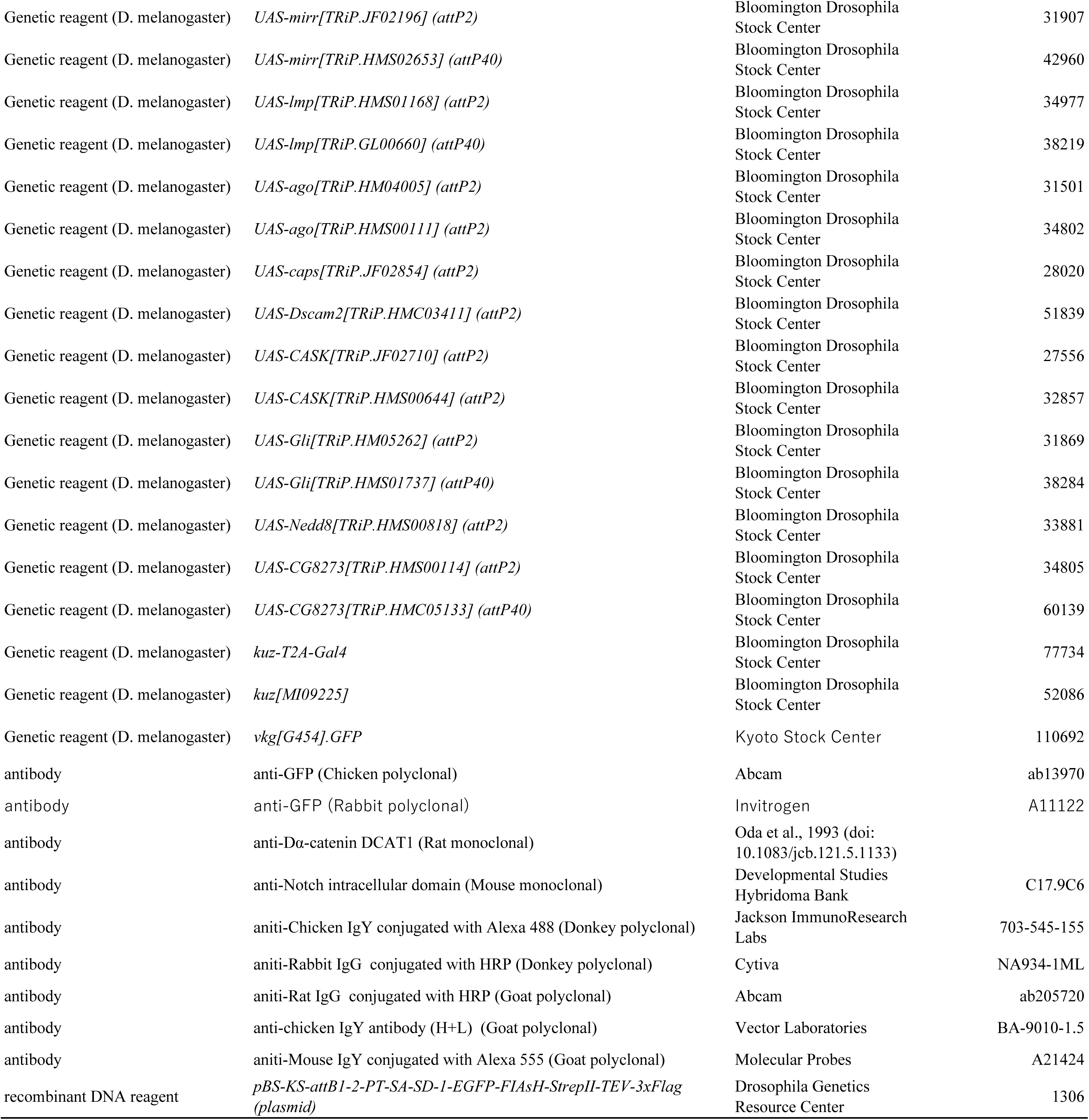
Fly stocks and reagents used in this study and exact genotypes and sex of flies in individual figure panels.

| Reagent type | Designation | Source | Identifiers |
| --- | --- | --- | --- |
| Genetic reagent (D. melanogaster) | *Canton-S* | EHIME-Fly Drosophila Stocks of Ehime University | E-10002 |
| Genetic reagent (D. melanogaster) | *w[1118]* |  |  |
| Genetic reagent (D. melanogaster) | *OK6-Gal4* | Sanyal, 2009 (doi: 10.1016/j.gep.2009.01.002) |  |
| Genetic reagent (D. melanogaster) | *c833-Gal4* | Bloomington Drosophila Stock Center | 6988 |
| Genetic reagent (D. melanogaster) | *ppl-Gal80 (attP2)* | Tsuyama et al., 2023 (doi: 10.1242/dev.200815) | *ppl-4kb-Gal80 (attP2)* |
| Genetic reagent (D. melanogaster) | *tub-Gal80[ts].20* | Bloomington Drosophila Stock Center | 7019 |
| Genetic reagent (D. melanogaster) | *elav-Gal80* | Simpson et al., 2016 (doi: 10.1080/01677063.2016.1248761) |  |
| Genetic reagent (D. melanogaster) | *UAS-CD4::tdGFP.VK00033* | Gift from Chun Han | PMID: 21606367 |
| Genetic reagent (D. melanogaster) | *UAS-mCD8::GFP* |  |  |
| Genetic reagent (D. melanogaster) | *UAS-lexA[TRiP.HMS05772] (attP40)* | Bloomington Drosophila Stock Center | 67947 |
| Genetic reagent (D. melanogaster) | *UAS-mCherry.VALIUM20 (attP2)* | Bloomington Drosophila Stock Center | 35785 |
| Genetic reagent (D. melanogaster) | *UAS-kuz[TRiP.HMS05424] (attP40)* | Bloomington Drosophila Stock Center |  |
| Genetic reagent (D. melanogaster) | *UAS-kuz[KK103555]* | Vienna Drosophila Resource Center | v107036 |
| Genetic reagent (D. melanogaster) | *UAS-kuz[VSH330255]* | Vienna Drosophila Resource Center | v330255 |
| Genetic reagent (D. melanogaster) | *UAS-srp[TRiP.HMS01298] (attP2)* | Bloomington Drosophila Stock Center | 34623 |
| Genetic reagent (D. melanogaster) | *UAS-N[TRiP.HMS00001] (attP2)* | Bloomington Drosophila Stock Center | 33611 |
| Genetic reagent (D. melanogaster) | *UAS-N[TRiP.JF01043] (attP2)* | Bloomington Drosophila Stock Center | 31502 |
| Genetic reagent (D. melanogaster) | *UAS-htl[TRiP.HMS04514] (attP40)* | Bloomington Drosophila Stock Center | 57313 |
| Genetic reagent (D. melanogaster) | *UAS-htl[TRiP.HMS01437] (attP2)* | Bloomington Drosophila Stock Center | 35024 |
| Genetic reagent (D. melanogaster) | *UAS-sr[TRiP.JF02781] (attP2)* | Bloomington Drosophila Stock Center | 27701 |
| Genetic reagent (D. melanogaster) | *UAS-st[TRiP.HMC05017] (attP2)* | Bloomington Drosophila Stock Center | 60024 |
| Genetic reagent (D. melanogaster) | *UAS-numb[TRiP.JF01695] (attP2)* | Bloomington Drosophila Stock Center | 31182 |
| Genetic reagent (D. melanogaster) | *UAS-numb[TRiP.HMS01459] (attP2)* | Bloomington Drosophila Stock Center | 35045 |
| Genetic reagent (D. melanogaster) | *UAS-haf[TRiP.HM05014] (attP2)* | Bloomington Drosophila Stock Center | 28528 |
| Genetic reagent (D. melanogaster) | *UAS-haf[TRiP.HMC03152] (attP2)* | Bloomington Drosophila Stock Center | 51438 |
| Genetic reagent (D. melanogaster) | *UAS-mirr[TRiP.JF02196] (attP2)* | Bloomington Drosophila Stock Center | 31907 |
| Genetic reagent (D. melanogaster) | *UAS-mirr[TRiP.HMS02653] (attP40)* | Bloomington Drosophila Stock Center | 42960 |
| Genetic reagent (D. melanogaster) | *UAS-lmp[TRiP.HMS01168] (attP2)* | Bloomington Drosophila Stock Center | 34977 |
| Genetic reagent (D. melanogaster) | *UAS-lmp[TRiP.GL00660] (attP40)* | Bloomington Drosophila Stock Center | 38219 |
| Genetic reagent (D. melanogaster) | *UAS-ago[TRiP.HM04005] (attP2)* | Bloomington Drosophila Stock Center | 31501 |
| Genetic reagent (D. melanogaster) | *UAS-ago[TRiP.HMS00111] (attP2)* | Bloomington Drosophila Stock Center | 34802 |
| Genetic reagent (D. melanogaster) | *UAS-caps[TRiP.JF02854] (attP2)* | Bloomington Drosophila Stock Center | 28020 |
| Genetic reagent (D. melanogaster) | *UAS-Dscam2[TRiP.HMC03411] (attP2)* | Bloomington Drosophila Stock Center | 51839 |
| Genetic reagent (D. melanogaster) | *UAS-CASK[TRiP.JF02710] (attP2)* | Bloomington Drosophila Stock Center | 27556 |
| Genetic reagent (D. melanogaster) | *UAS-CASK[TRiP.HMS00644] (attP2)* | Bloomington Drosophila Stock Center | 32857 |
| Genetic reagent (D. melanogaster) | *UAS-Gli[TRiP.HM05262] (attP2)* | Bloomington Drosophila Stock Center | 31869 |
| Genetic reagent (D. melanogaster) | *UAS-Gli[TRiP.HMS01737] (attP40)* | Bloomington Drosophila Stock Center | 38284 |
| Genetic reagent (D. melanogaster) | *UAS-Nedd8[TRiP.HMS00818] (attP2)* | Bloomington Drosophila Stock Center | 33881 |
| Genetic reagent (D. melanogaster) | *UAS-CG8273[TRiP.HMS00114] (attP2)* | Bloomington Drosophila Stock Center | 34805 |
| Genetic reagent (D. melanogaster) | *UAS-CG8273[TRiP.HMC05133] (attP40)* | Bloomington Drosophila Stock Center | 60139 |
| Genetic reagent (D. melanogaster) | *kuz-T2A-Gal4* | Bloomington Drosophila Stock Center | 77734 |
| Genetic reagent (D. melanogaster) | *kuz[MI09225]* | Bloomington Drosophila Stock Center | 52086 |
| Genetic reagent (D. melanogaster) | *vkg[G454].GFP* | Kyoto Stock Center | 110692 |
| antibody | anti-GFP (Chicken polyclonal) | Abcam | ab13970 |
| antibody | anti-GFP (Rabbit polyclonal) | Invitrogen | A11122 |
| antibody | anti-Dα-catenin DCAT1 (Rat monoclonal) | Oda et al., 1993 (doi: 10.1083/jcb.121.5.1133) | |
| antibody | anti-Notch intracellular domain (Mouse monoclonal) | Developmental Studies Hybridoma Bank | C17.9C6 |
| antibody | aniti-Chicken IgY conjugated with Alexa 488 (Donkey polyclonal) | Jackson ImmunoResearch Labs | 703-545-155 |
| antibody | aniti-Rabbit IgG conjugated with HRP (Donkey polyclonal) | Cytiva | NA934-1ML |
| antibody | aniti-Rat IgG conjugated with HRP (Goat polyclonal) | Abcam | ab205720 |
| antibody | anti-chicken IgY antibody (H+L) (Goat polyclonal) | Vector Laboratories | BA-9010-1.5 |
| antibody | aniti-Mouse IgY conjugated with Alexa 555 (Goat polyclonal) | Molecular Probes | A21424 |
| recombinant DNA reagent | *pBS-KS-attB1-2-PT-SA-SD-1-EGFP-FIAsH-StrepII-TEV-3xFlag (plasmid)* | Drosophila Genetics Resource Center | 1306 |

**Table S3.**
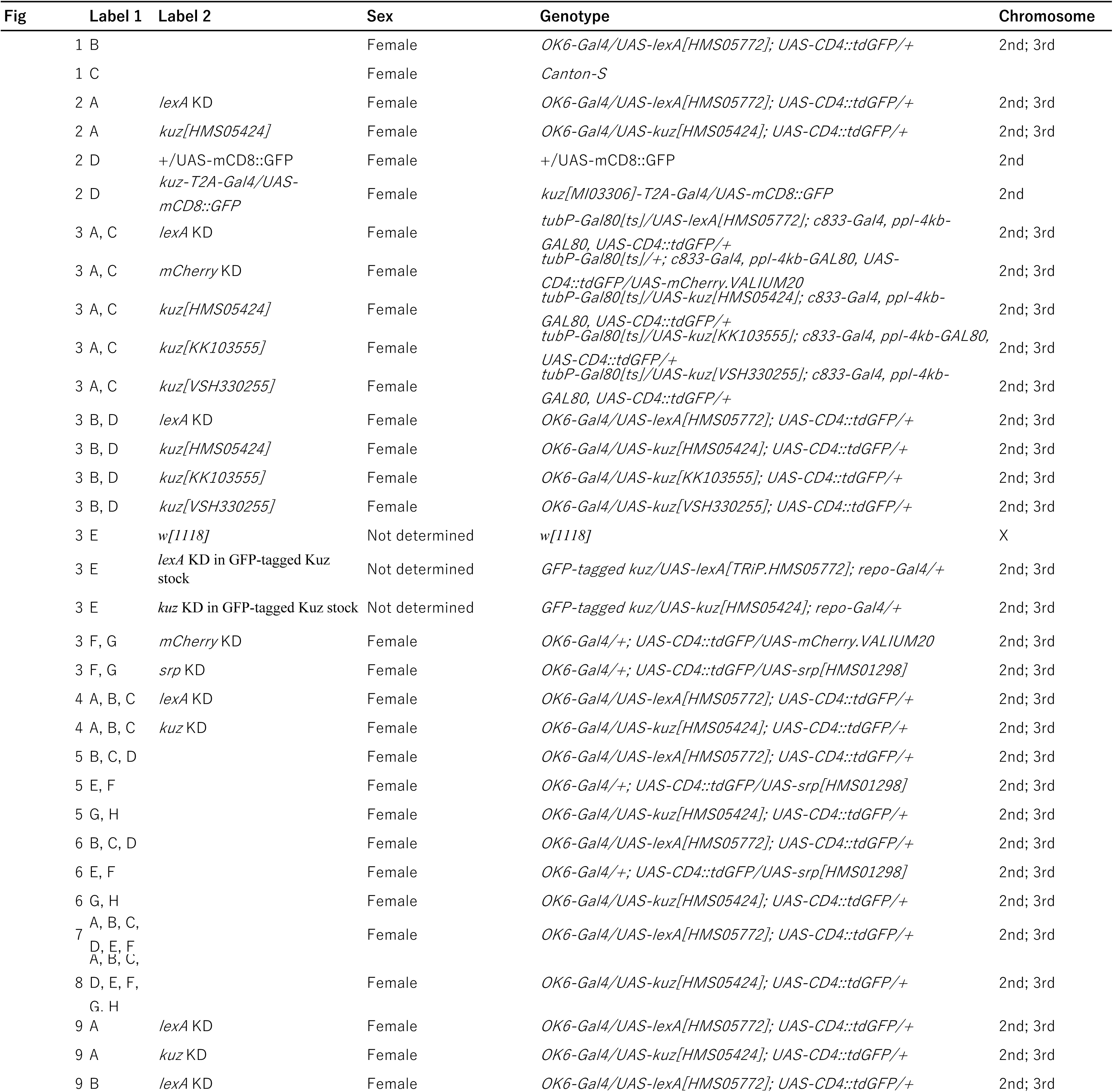
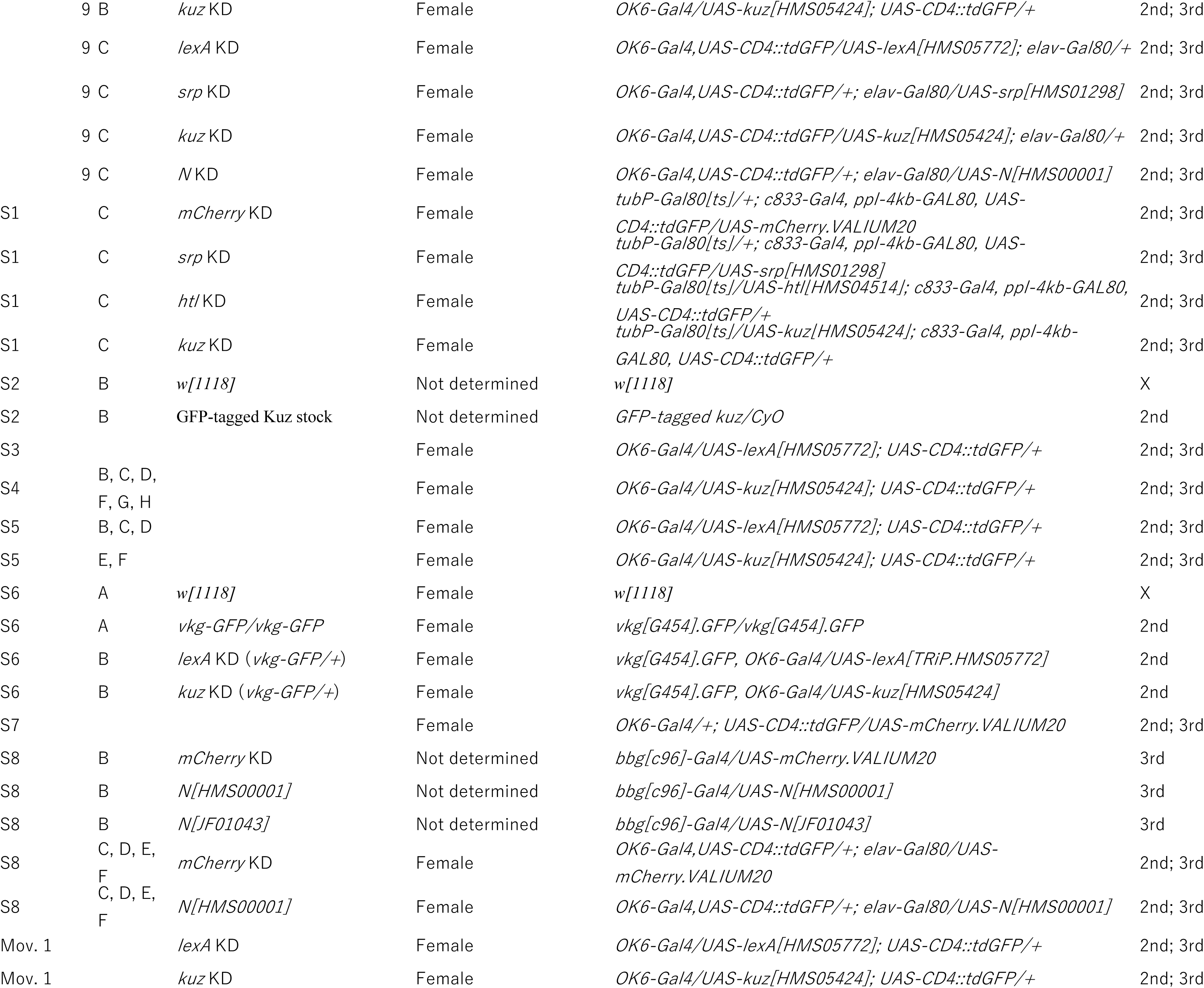
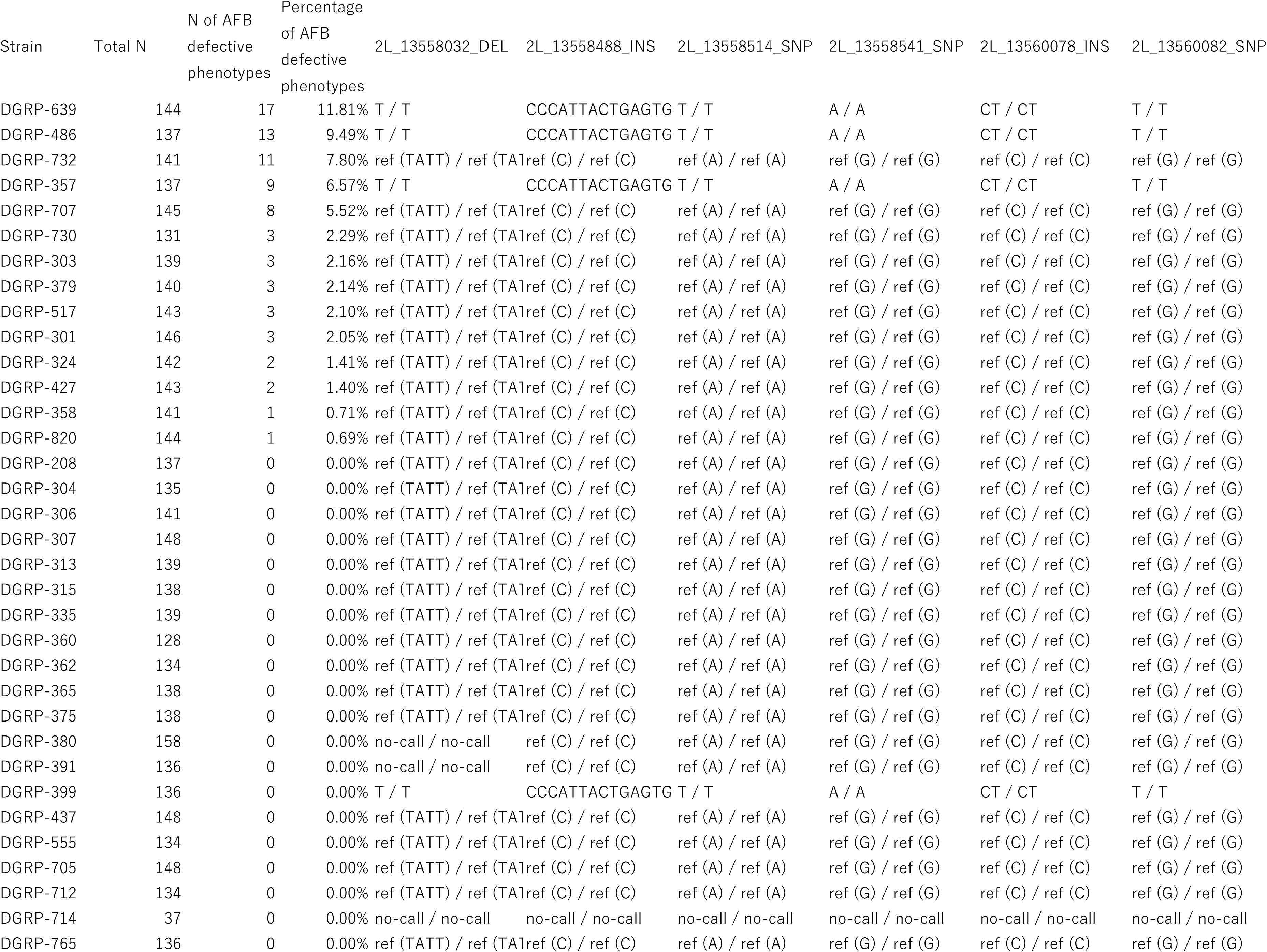
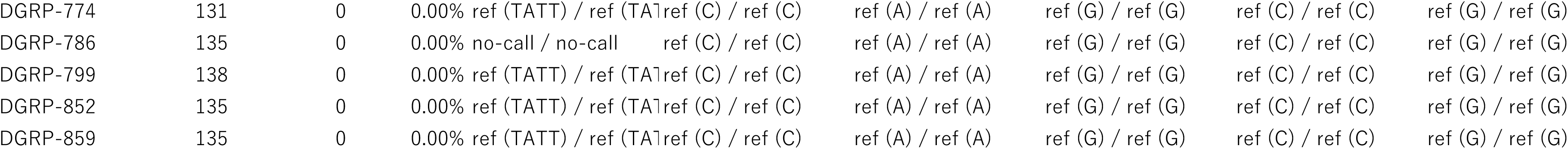
Genetic variants of *kuz* in the representative 39 strains in *Drosophila* Genetic Reference Panel.

## Notes

### Competing Interest Statement

The authors have declared no competing interest.

